# Connectivity establishes spatial readout of visual looming in a glomerulus lacking retinotopy

**DOI:** 10.1101/2020.04.19.036947

**Authors:** Mai M. Morimoto, Aljoscha Nern, Arthur Zhao, Edward M. Rogers, Allan M. Wong, Mathew D. Isaacson, Davi D. Bock, Gerald M. Rubin, Michael B. Reiser

## Abstract

Visual systems can exploit spatial correlations in the visual scene by using retinotopy, the organizing principle by which neighboring cells encode neighboring spatial locations. However, retinotopy is often lost, such as when visual pathways are integrated with other sensory modalities. How is spatial information processed in the absence of retinotopy? Here, we focused on visual looming responsive LC6 cells in *Drosophila*, a population whose dendrites collectively tile the visual field, but whose axons form a single glomerulus—a structure lacking retinotopic organization—in the central brain. We identified multiple glomerulus neurons and found that they respond to looming in different portions of the visual field, unexpectedly preserving spatial information. Through EM reconstruction of all LC6 synaptic inputs to the glomerulus, we found that LC6 and downstream cell types form circuits within the glomerulus that establish spatial readout of visual features and contralateral suppression—mechanisms that transform visual information for behavioral control.

## Introduction

In many animals, brain regions involved in processing visual information are large and well organized, featuring retinotopy—an organizational plan that preserves the mapping of space originating in the retina, such that neighboring neurons respond to visual signals at neighboring spatial locations. Animals need visual-spatial information in order to direct their escape away from predators, to find mates, or to catch prey. Classical studies in cats (Hubel and Wiesel, 1962) and monkeys (Tootell et al., 1988), and recent work in mice (Garrett et al., 2014) demonstrated that the retinotopic organization of higher visual areas is a dominant feature of the organization of the mammalian brain. While this organizing principle has facilitated detailed analyses of visual pathways, it remains unclear how visual-spatial information is processed in the absence of retinotopy. In particular, retinotopy must be sacrificed where visual information is integrated with other modalities and as vision is translated into behavioral actions. We propose that rapid progress can be made on understanding these critical transformations in the *Drosophila* brain.

The lobula columnar (LC) neurons project from the visual system into the central brain of flies, and have been well described in *Drosophila* (Fischbach and Dittrich, 1989; Otsuna and Ito, 2006; Wu et al., 2016). They are positioned at the last processing step of a retinotopic neuropil and are a nexus between the detection of visual features and the organization of behavioral control. There are ∼20 types of LC cells, and we have previously shown that optogenetic activation of individual cell types elicits a wide range of behaviors that closely resemble natural behaviors such as escape jumps, backward walking, courtship behavior, and reaching (Wu et al., 2016). Furthermore, the cell types that elicited avoidance behaviors (LC4, LPLC2, LC6 and LC16) also responded to visual looming stimuli (Ache et al., 2019; Klapoetke et al., 2017; Sen et al., 2017; Wu et al., 2016). Other cell types, including cells whose activation elicited courtship or reaching behaviors (LC10), responded to the motion of small visual objects (Keleş and Frye, 2017; Ribeiro et al., 2018; Wu et al., 2016). These findings suggest that LC neurons encode ethologically relevant visual object features and that these are translated in to appropriate visually guided actions by downstream circuits.

Anatomical studies, based on light microscopy, have suggested that most LC neurons (with the notable exception of LC10) lose retinotopy—their inputs in the visual system are clearly organized with the spatial layout of the retina, while their axonal projections, into glomeruli in the central brain, appear to lack any continuous visual-spatial organization (Mu et al., 2012; Wu et al., 2016). This anatomical evidence has led to the suggestion that LC neurons may represent a critical transition from visual (‘where’) processing to a more abstract (‘what’) representation of visual information (Mu et al., 2012; Strausfeld et al., 2007; Wu et al., 2016).

In this study we have systematically explored the downstream circuitry of one LC type. We focused on LC6 (Figure 1A-B), a cell type that elicited the most reliable escape take-off behavior when depolarized (optogenetic activation screen; (Wu et al., 2016)), but is not a direct input to the well-studied giant-fiber escape pathway (Ache et al., 2019). The LC6 axons are presynaptic within the LC6 glomerulus, a structure that does not exhibit retinotopic organization (Figure 1C, C’, D). We identified several neurons with arbors in the LC6 glomerulus and established genetic tools for targeting these cell types. We assayed the functional connectivity between LC6 neurons and these candidate downstream cell types, yielding five distinct cell types that integrate LC6 inputs. We further investigated two of these downstream neuron types, one that connected the two LC6 glomeruli across both hemispheres, and an ipsilateral type that projected to a higher order multi-sensory area (anterior ventrolateral protocerebrum, AVLP). Using calcium imaging we examined how these downstream targets further modified the stimulus selectivity of LC6 neurons. During this analysis we were surprised to find that LC6 target neurons read out spatially selective information from the LC6 glomerulus, despite the lack of retinotopic organization in this structure.

**Figure 1:**
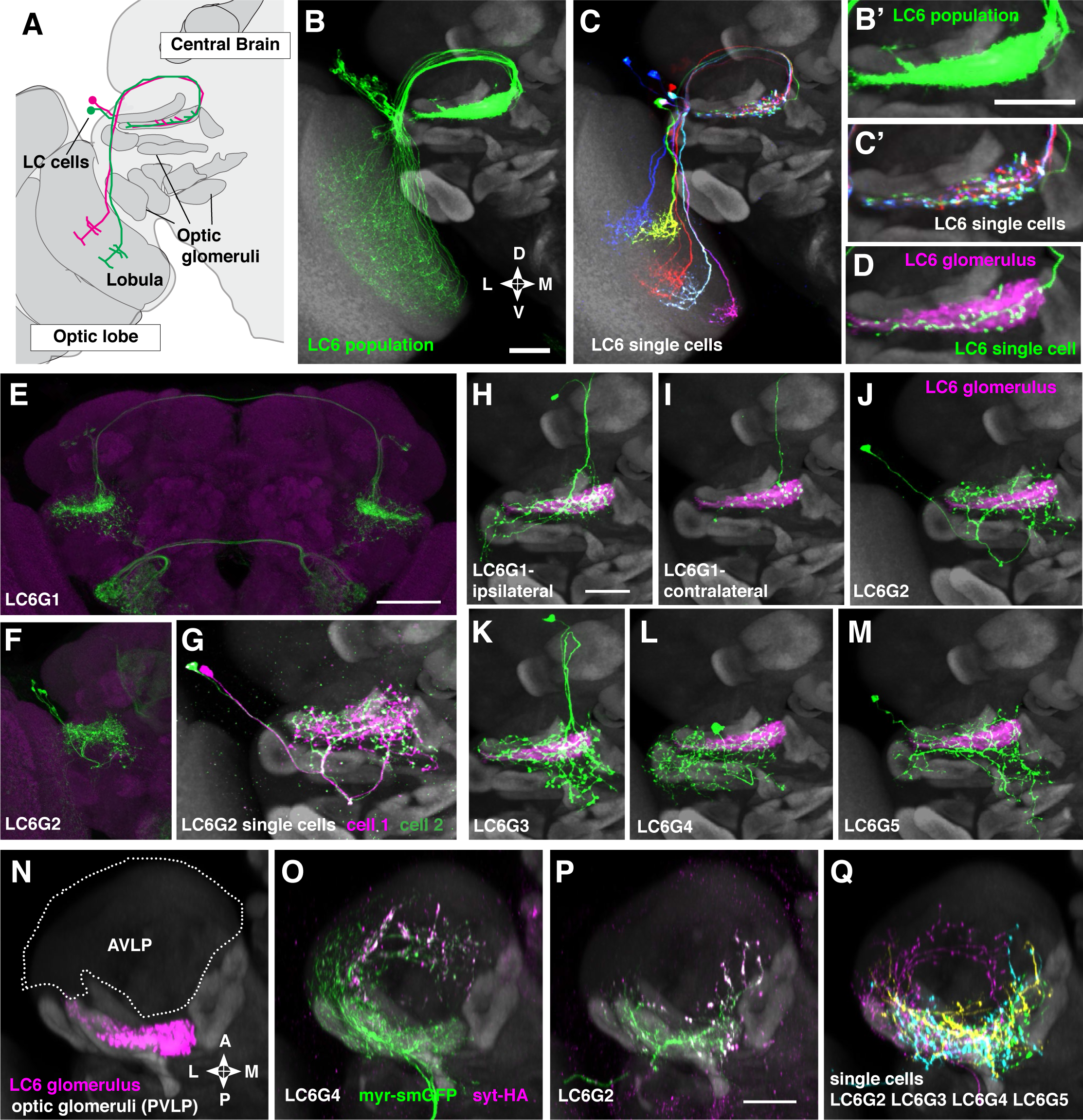
Multiple distinct cell types innervate the LC6 glomerulus. **(A-D)** Projection pattern of LC6 neurons. **(A)** Lobula Columnar (LC) neurons project from the lobula to synapse rich structures in the posterior ventrolateral protocerebrum (PVLP) called optic glomeruli. Two LC6 cells projecting to the LC6 glomerulus are illustrated. Adapted from (Wu et al 2016). In all panels except for N-Q dorsal is up and medial is to the right. **(B, C)** LC6 neuron dendrites collectively tile the lobula, while the axons converge to form a glomerulus. **(B, B’)** Population of LC6 neurons labeled with a membrane targeted marker. All panels (except **E**, **F**) show composites of registered confocal images together with the standard brain used for registration (shown in grey). To more clearly show the neurons of interest, some images were manually segmented to exclude additional labeled cells or background signal (see Methods). Scale bar, 20 µm. **(C, C’)** MultiColor FlpOut (MCFO)-labeled individual LC6 cells registered and displayed as in **B**. Note that individual LC6 terminals overlap in the LC6 glomerulus, while dendrites in the lobula occupy distinct positions. **(D)** Terminal of a single MCFO-labeled LC6 cell displayed as in **B’** with the LC6 glomerulus, based on the expression of a synaptic marker (syt-HA) in LC6 cells as described in (Wu et al 2016), in magenta. **(E, F)** Examples of two populations of candidate LC6 target cells labeled by split-GAL4 driver lines. A membrane marker is in green, a synaptic marker (anti-Brp) in magenta. Scale bar, 50 µm. **(G)** MCFO labeling of two LC6G2 cells in the same specimen. **(H-M)** Examples of single cells of potential LC6 targets. All images show MCFO-labeled manually segmented single cells (green) together with the standard brain (grey) and LC6 glomerulus (magenta). Scale bar, 20 µm. **(N-O)** Several LC6 glomerulus interneurons project to the anterior ventrolateral protocerebrum (AVLP). The images show projections through a sub stack rotated around the mediolateral axis by 90° relative to the view shown in the other panels; reference brain neuropil marker (grey), LC6 glomerulus (magenta) and approximate boundaries of the AVLP (dotted white line). **(O, P)** LC6G4 and LC6G2 populations, as labeled by split-GAL4 lines. In addition to the membrane label (green) and standard brain (grey) a presynaptic marker (syt-HA) is shown (magenta). Note syt-HA labeling in the AVLP. **(Q)** Overlay of the segmented single cells of LC6G2, LC6G3, LC6G4, and LC6G5 shown in **J-M** with the standard brain. Images in **N-Q** are sub stack projection, some cells in **O-Q** have additional AVLP branches outside of the projected volume. See Table 1 for detailed summary of fly lines used in this study.

**Table 1:** Detailed summary of genotypes used throughout the manuscript.

| Figure | Driver line(s) | Reporter / effector transgene(s) | Description |
| --- | --- | --- | --- |
| 1B, 1B' | OL0070B | pJFRC51-3XUAS-IVS-syt::smHA (su(Hw)attP1) ,pJFRC225-5XUAS-IVS-myr::smFLAG (VK00005) (only myr::smFLAG pattern shown) | Composite image includes reference brain pattern<br>63x objective |
| 1C, 1C' | OL0081B | MCFO-1 [pBPhsFlp2::PEST (attP3);; pJFRC201- 10XUAS-FRT>STOP>FRT-myr::smGFP-HA (VK0005), pJFRC240-10XUAS-FRT> STOP>FRT-myr::smGFP-V5-THS-10XUASFRT>STOP>FRT-myr::smGFP-FLAG (su(Hw)attP1)] | Composite image includes reference brain pattern<br>(63x objective) |
| 1D,1H,1I, 1J,1K, 1L,1M, 1N, 1—fs1 | OL0070B | pJFRC51-3XUAS-IVS-syt::smHA (su(Hw)attP1) ,pJFRC225-5XUAS-IVS-myr::smFLAG (VK00005) (only syt::smHA pattern shown) | LC6 glomerulus label used in several panels (manually segmented)<br>(63x objective) |
| 1D | OL0081B (single cell) | MCFO-1 | Composite image includes reference brain pattern and LC6 glomerulus label<br>(63x objective) |
| 1E | SS00825 | pJFRC51-3XUAS-IVS-syt::smHA (su(Hw)attP1) ,pJFRC225-5XUAS-IVS-myr::smFLAG (VK00005) (only myr::smFLAG pattern shown)<br>anti-Brp neuropile marker | Maximum intensity projection<br>(20x objective) |
| 1F | SS02036 | pJFRC51-3XUAS-IVS-syt::smHA (su(Hw)attP1) ,pJFRC225-5XUAS-IVS-myr::smFLAG (VK00005) (only myr::smFLAG pattern shown)<br>anti-Brp neuropile marker | Maximum intensity projection<br>(20x objective) |
| 1G | SS02036 | MCFO-1 | Composite images include reference brain pattern and LC6 glomerulus label<br>Manually segmented cells.<br>(63x objective) |
| 1H, 1—fs1A | SS00825 | MCFO-1 |  |
| 1I, 1—fs1B | SS00824 | MCFO-1 |  |
| 1J, 1—fs1C | SS02036 | MCFO-1 |  |
| 1K, 1—fs1D | SS02699 | MCFO-1 |  |
| 1L, 1—fs1E | SS03690 | MCFO-1 |  |
| 1M, 1—fs1F | SS02410 | MCFO-1 |  |
| 1N | NA | NA | Composite image of reference brain pattern and LC6 glomerulus label<br>(63x objective) |
| 1O | SS03690 | pJFRC51-3XUAS-IVS-syt::smHA (su(Hw)attP1) ,pJFRC225-5XUAS-IVS-myr::smFLAG (VK00005) | Composite image with reference brain pattern<br>(63x objective) |
| 1P | SS02036 | pJFRC51-3XUAS-IVS-syt::smHA (su(Hw)attP1) ,pJFRC225-5XUAS-IVS-myr::smFLAG (VK00005) | Composite image with reference brain pattern (63x objective) |
| 1Q | SS02036, SS02699, SS03690, SS02410 | MCFO-1 | Composite image with reference brain pattern<br>Overlay of cells from 1J, 1K, 1L, 1M (shown in a different view) |
| 2, 2—fs1: LC6 | OL0070B | FC-1 [13XLexAop2-CsChrimson-tdT (attP18), 20XUAS-IVS-Syn21-opGCaMP6f p10 Su(Hw)attP8); 42E06-LexA(JK22C);+] | Functional connectivity experiments: depolarizing LC6 with Chrimson, while imaging calcium responses from other cell type. Used with two-photon imaging. |
| 2, 2—fs1: LC6G1 | SS00825 | FC-1 |  |
| 2, 2—fs1: LC6G2 | SS02036 | FC-1 |  |
| 2, 2—fs1: LC6G3 | SS02099 | FC-1 |  |
| 2, 2—fs1: LC6G4 | SS03690 | FC-1 |  |
| 2, 2—fs1: LC6G5 | SS02409 | FC-1 |  |
| 2—fs1: LC26G1 | SS03641 | FC-1 |  |
| 3, 4: LC6 | OL0070B | GCaMP-1 [pJFRC7-20XUAS-IVS-GCaMP6m (VK00005)] | Calcium imaging from either LC6 or one of the LC6G neurons. In a subset of the recordings, the LC6 glomerulus was labeled with tdTomato. Used with two-photon imaging. |
| 3, 4, 3—fs1: LC6G1 | SS00825 | GCaMP-1 or GCaMP-2 [pJFRC48-13XLexAop2-IVS-myrtdTomato (su(Hw)attP8); 42E06-LexA(JK22C); pJFRC7-20XUAS-IVS-GCaMP6m (VK00005)] |  |
| 3, 4, 3—fs1, 7: LC6G2 | SS02036 | GCaMP-1 or GCaMP-2 |  |
| 1—fs1G, 1—fs1G', 1—fs1H, 1—fs1H' | SS03641 | pJFRC51-3XUAS-IVS-syt::smHA (su(Hw)attP1) ,pJFRC225-5XUAS-IVS-myr::smFLAG (VK00005) | Composite image with reference brain pattern and LC26 glomerulus label based on syt-HA expression. Manually segmented. (63x objective) |
| 1—fs2 | SS00825, SS02036, SS02099, SS02409, SS03690, SS03641 | pJFRC51-3XUAS-IVS-syt::smHA (su(Hw)attP1) ,pJFRC225-5XUAS-IVS-myr::smFLAG (VK00005) (only myr::smFLAG pattern shown)<br>anti-Brp neuropile marker | Maximum intensity projections<br>(20x objective) |
| 1—fs3 | SS00825, SS02036, SS02099, SS03690, SS25111 | *UAS-7xHaloTag::CAAX* in *VK00005*<br>FISH probes for GAD1, VGluT, ChAT mRNAs | Single confocal sections<br>(63x objective) |

fs = figure supplement

What if visual-spatial information is conveyed not by the innervation pattern of these projections, but rather by specific synaptic connections? To determine where this specificity for spatial information originates, we undertook a comprehensive analysis of the connectivity between LC6 neurons and two downstream cell types using recently available whole-brain Electron Microscopy (EM) data (Zheng et al., 2018). We reconstructed anatomical ‘receptive fields’ for all LC6 neurons and used them to estimate the receptive fields of the connected downstream cells. We found that downstream neurons access distinct spatial information via biased connections with LC6s, consistent with our functional measurements. Finally, combining connectivity and functional data, we detail a circuit wherein a downstream target of LC6 receives contralateral suppression that enhances the detection of looming stimuli over confounding visual cues.

## Results

### Multiple distinct cell types innervate the LC6 glomerulus

In *Drosophila* it has become increasingly feasible to identify genetic markers for cell types of interest by analyzing images of GAL4 line expression patterns (Jenett et al., 2012a; Kvon et al., 2014; Panser et al., 2016). To find potential LC6 targets, we visually searched for neurons with processes that substantially overlap with the LC6 glomerulus, a distinct neuropil structure that contains the densely packed terminals of LC6 neurons (Figure 1A-D). This approach identified several candidate LC6 glomerulus interneurons (Figure 1—figure supplement 1). To facilitate studies of these cells, we developed more specific split-GAL4 driver lines (Luan et al., 2006; Pfeiffer et al., 2010) (Figure 1E, F, Figure 1—figure supplement 2). We obtained split-GAL4 lines with expression in each of five distinct cell populations that heavily overlap with the LC6 glomerulus. We named these putative cell types, LC6G1 - LC6G5 (short for LC6 Glomerulus 1-5; Figure 1H-M, Figure 1—figure supplement 1). LC6G types have identifying characteristic cell body locations and projection patterns (Figure 1E-F, Figure 1—figure supplement 1, 2). Cell body positions, presumably reflecting distinct developmental origins, are in the cell body rind of the anterior dorsal (LC6G1 and LC6G3) or posterior (LC6G4) central brain and in the space between optic lobe and central brain (LC6G2 and LC6G5). Bilateral arbors that span both brain hemispheres differentiate LC6G1 from the other LC6Gs (which have only ipsilateral processes; Figure 1E, F, Figure 1—figure supplement 2). Expression of neurotransmitter markers (VGlut, GAD1, ChaT) also indicates differences between LC6G types (Figure 1—figure supplement 3): LC6G1 and LC6G3 appear to be glutamatergic, LC6G4 GABAergic, and LC6G2 and LC6G5 cholinergic. In the fly CNS, the expression of a glutamate-gated chloride channel indicates that glutamate can act as an inhibitory neurotransmitter (Liu and Wilson, 2013; Mauss et al., 2015; Strother et al., 2017), suggesting that the LC6G1, LC6G3, and LC6G4 neurons may provide inhibition of their targets in the LC6 glomerulus. On the other hand, LC6G2, LC6G5, and LC6 neurons express ChaT (Figure 1—figure supplement 3 and (Davis et al., 2020)) and so are likely to be cholinergic, and thus excitatory neurons. Single-cell labeling revealed further details of each group (Figure 1H-M, G, Figure 1—figure supplement 1). While we focus here on the LC6 glomerulus, our search also identified neurons with arbors in other glomeruli. One example, a potential LC26 target (LC26G1) is shown in Figure 1—figure supplement 1; we use this cell type as a control in one of our experiments.

All LC6G cells have at least some branches outside of the glomerulus. In several cases, these extend into other glomeruli, including, for example, the LC16 target region in the case of LCG6G2 (Figure 1J) and the LC15 and LC21 glomeruli for LC6G4 (Figure 1L). Apart from other glomeruli and their immediate surroundings, the AVLP (Anterior Ventrolateral Protocerebrum) appears to be the main target region for ipsilateral LC6G cells (Figure 1N-Q). In particular, LC6G2, LC6G4, LC6G5 have potential presynaptic sites in this region, which is presumed to be a higher order, multi-sensory area and contains both arbors of descending interneurons (Namiki et al., 2018) and interneurons projecting to other central brain regions. Thus, these LC6G cells have the anatomy of projection neurons, not local neurons.

Each LC6G split-GAL4 driver labels multiple individual cells (disregarding unrelated cells in distinct regions also present in some lines). For several driver lines, we used multicolor stochastic labeling (Nern et al., 2015) to directly visualize multiple similar cells in the same specimen (Figure 1G, Figure 1—figure supplement 1). There are about 4-6 LC6G1, LC6G2, LC6G3 and LC6G4 cells each per hemisphere. While these numbers could include a few unrecognized cells of other types or underestimate the true cell number due to incomplete driver expression, each LC6G type appears to consist of multiple cells of very similar or identical morphology. Similar to LC6 cells (Figure 1C, D), the distribution of processes of LC6G cells does not subdivide the LC6 glomerulus into distinct subregions. If these neurons are indeed LC6 targets, it is unclear whether they receive input from most or all LC6 cells or from random or specific subsets.

### Pairwise functional connectivity reveals connected downstream cell-types

To assess whether the candidate downstream neuron types identified by our anatomical analysis are functionally connected to LC6s, we optogenetically depolarized LC6 neurons and measured the calcium activity of each candidate downstream cell type. In a series of pairwise experiments, the candidate downstream split-GAL4 lines drove expression of GCaMP6f (T.-W. Chen et al., 2013), while the light-gated ion channel Chrimson (tagged with tdTomato; (Klapoetke et al., 2017; Strother et al., 2017)) was expressed in LC6 using the orthogonal LexA/LexAop system (Lai and Lee, 2006; Pfeiffer et al., 2010) (Figure 2A, left, LC6 in magenta, candidate downstream LC6Gs in green). Flies were dissected and whole brains (not including the photoreceptors) were imaged using two-photon microscopy (Figure 2A, right). To obtain an LC6 activity-dependent tuning curve for responses of downstream neurons, we first conducted a “calibration” experiment to select a series of increasing light stimulation intensities that would evoke activity in LC6s with monotonically increasing responses. The stimulation light was spatially restricted to the glomerulus being imaged (detailed in methods). We established a six-pulse stimulation protocol with ramping light intensity with which we could observe a monotonically increasing LC6 GCaMP signal in response to increasing levels of light-evoked Chrimson activation (Figure 2C, D; LC6).

**Figure 2:**
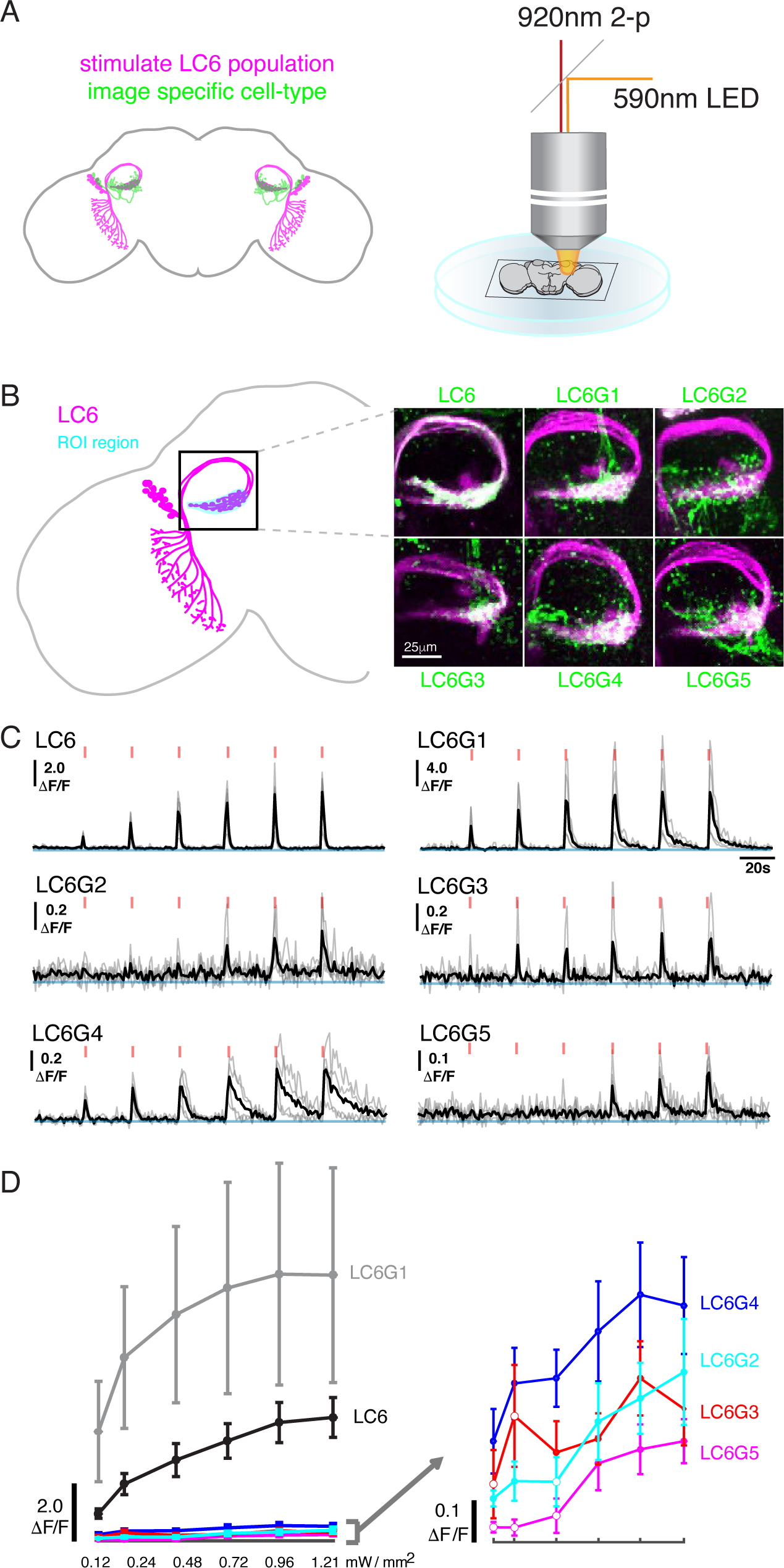
Pairwise functional connectivity reveals connected downstream cell-types. **(A)** Chrimson was expressed in LC6 (magenta) using a LexA line while GCaMP6f was expressed in individual candidate downstream neuron types (green) using split-GAL4 lines. Chrimson was activated by pulses of 590 nm light while GCaMP responses were imaged using two-photon microscopy. **(B)** Calcium responses were measured in the ROI region in cyan. Zoom-in of the LC6 glomerulus region show representative images of double-labeled (Chrimson in magenta, GcaMP6f in green) brains used for the experiment. **(C)** Calcium responses in candidate downstream neurons in response to LC6 Chrimson activation (N=4-5 per combination; individual sample response in gray, mean response in black). A range of response amplitudes and timecourses were observed. Red tick marks indicate the activation stimulus. **(D)** Peak responses (mean ± SEM) are shown for each candidate downstream cell type for increasing light stimulation. Closed circles denote data points significantly different from pre-stimulus baseline, open circles denote data points not significantly different from pre-stimulus baseline (p<0.1, two-sample t-test). See Table 1 for detailed summary of fly lines used in this study.

Using this stimulation protocol, we measured responses from the candidate downstream neuron types while activating LC6 neurons (Region of Interest, ROI, Figure 2B, in cyan). All cell types, except LC26G1 (control line), showed increasing calcium responses to LC6 activation (alternative stimulation protocol and control line responses shown in Figure 2— figure supplement 1). The responses of the different downstream neuron types had different amplitudes and temporal kinetics (Figure 2C, Figure 2—figure supplement 1A), likely reflecting a combination of connection strengths to LC6, intrinsic properties of the target neurons, indirect contributions evoked by the stimulation, and possible expression strength differences. The amplitude difference is most clear in the response curves (Figure 2D, Figure 2—figure supplement 1B): the bilateral LC6G1 neurons showed a strikingly large ΔF/F (∼9.0), whereas the other cell types showed an order of magnitude smaller increase in ΔF/F (∼0.4). Part of this difference is due to the density of processes of each cell type. The LC6G1 neurons fill the glomerulus more densely than the other LC6Gs. However, this difference is unlikely to account for a ∼20X difference in response magnitude. All cells proposed to be downstream of LC6 (Figure 1) showed significant responses (results of statistical analysis in Figure 2D, 2—figure supplement 1B) to strong levels of LC6 activation, consistent with their being directly postsynaptic to LC6 axons in the glomerulus.

### Neurons downstream of LC6 exhibit stereotyped spatial receptive fields

Having established multiple cell types as connected to LC6, we next examined whether these cells respond to the same visual looming stimuli that selectively activate LC6 neurons (Wu et al., 2016). For further analysis we selected the strongly connected, bilaterally projecting LC6G1s as well as the LC6G2 ipsilaterally projecting neurons. We expressed GCaMP6m (T. Chen et al., 2013) using the same split-GAL4 driver lines used for the targeted functional connectivity experiments, and measured calcium responses from the LC6Gs (summed response from ∼four cells per brain hemisphere in each cell type) to visual stimuli using *in vivo* two-photon microscopy (Figure 3A, left). As with LC6 neurons, both LC6G types responded to visual looming stimuli (example response for LC6G2 recordings in Figure 3A, middle). However, the initial measurements appeared to show substantial selectivity for stimuli presented in different spatial locations. In light of the anatomy of the glomerulus, this spatial selectivity was unexpected. To confirm this, we mapped the responses to small looming disks (maximum size 18°) presented at all locations on a grid covering a large region of the right eye’s field of view. The resulting spatial Receptive Field (RF) is shown in Figure 3A (right). We precisely measured the head orientation (Figure 3—figure supplement 1A) to map the visual stimulus onto the fly’s eye (Figure 3—figure supplement 1B).

**Figure 3:**
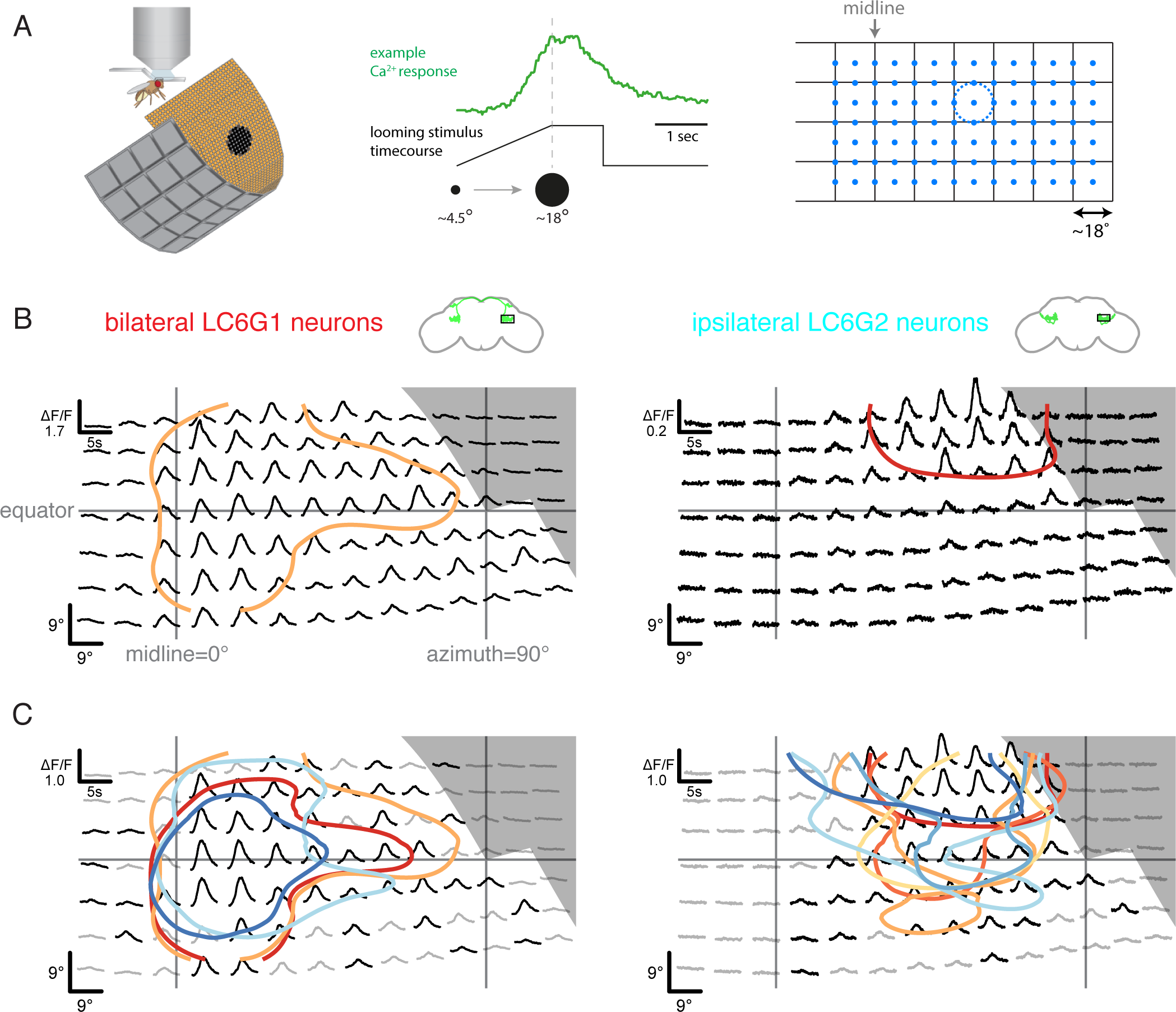
Neurons downstream of LC6 exhibit stereotyped spatial receptive fields. **(A)** Left: Schematic of the *in vivo* two-photon calcium imaging setup. An LED display was used to deliver looming stimuli at many positions around the fly, while calcium responses were measured from an ROI containing the LC6 glomerulus. Middle: Example stimulus time course and response of an LC6G2 neuron. Right: Small looming stimuli were delivered at 98 positions centered on a grid (and subtending ∼18° diameter at maximum size) to map the receptive fields of LC6G neurons. **(B, C)** The bilateral LC6G1 and ipsilateral LC6G2 neurons showed consistent visual responses within restricted receptive fields. **(B)** Single animal example responses, with each time series showing the mean of 3 repetitions of the stimulus at each location. The responses are plotted to represent the grid of looming stimuli mapped onto fly eye centered spherical coordinates (see Figure 3—figure supplement 1B; lines indicating the eye’s equator, frontal midline, and lateral 90°, are shown in gray). The (0,0) origin of each time series is plotted at the spatial position of the center of each looming stimulus (this alignment onto eye coordinates accounts for e.g. the upward curving stimulus responses below the eye equator). The 60% of peak response is demarcated with a contour (based on interpolated mean responses at each location). **(C)** The mean responses across multiple animals are shown (N=4 for LC6G1, N=7 for LC6G2), with the 60% of peak contours shown in a separate color for each individual fly. The responses have been normalized (to a peak response of ∼1) for each fly before averaging. Responses at each position that are significantly larger than zero are in black, others in gray (one-sample t-test, p<0.05, controlled for False Discovery Rate (Benjamini and Hochberg, 1995)). The individual responses are shown in Figure 3—figure supplement 1; the gray area indicates the region of the visual display that was occluded by the head mount (Figure 3—figure supplement 1B). See Table 1 for detailed summary of fly lines used in this study and Table 2 for summary of all visual stimuli presented.

**Table 2:** Detailed summary of all visual stimuli used throughout the manuscript.

| Stimulus type | Stimulus name | Nominal location (azimuth, elevation) | Speeds | Fig.3 | Fig.4B | Fig.4C | Fig.7B |
| --- | --- | --- | --- | --- | --- | --- | --- |
| 1 | Small looming RF mapping stimulus | 98 center positions ranging -18° to 99°, ±27° | 10°/s | small looming |  |  |  |
| 2 | Dark Looming Disc side | centered at 45°, -9° (18° for LC6G2) | r/v = 10, 40, 70, 130, 310, 550 ms |  | looming | looming 1 |  |
| 2 | Dark Receding Disc side | centered at 45°, -9° (18° for LC6G2) | r/v = 10, 40, 70, 130, 310, 550 ms |  | receding | receding 3 |  |
| 2 | Dark Looming Disc front | centered at 27°, -9° (18° for LC6G2) | r/v = 10, 40, 70, 130, 310, 550 ms |  |  | looming 4 |  |
| 3 | Dark Receding Disc front | centered at 27°, -9° (18° for LC6G2) | r/v = 10, 40, 70, 130, 310, 550 ms |  |  | receding 2 |  |
| 3 | Bright Looming Disc side | centered at 45°, -9° (18° for LC6G2) | r/v = 10, 40, 70, 130, 310, 550 ms |  | bright looming | looming 2 |  |
| 3 | Bright Receding Disc side | centered at 45°, -9° (18° for LC6G2) | r/v = 10, 40, 70, 130, 310, 550 ms |  |  | receding 1 |  |
| 4 | Dark Constant Looming Disc side | centered at 45°, -9° (18° for LC6G2) | 5,7,10,20,30,40,50,70,101,203,494°/s |  |  | looming 3 | ipsi, contra-only expanding |
| 4 | Dark Constant Receding Disc side | centered at 45°, -9° (18° for LC6G2) | 5,7,10,20,30,40,50,70,101,203,494°/s |  |  | receding 4 | ipsi-only receding |
| 5 | Luminance-matched stimulus side | centered at 45°, -9° (18° for LC6G2) | r/v = 10, 40, 70, 130, 310, 550 ms |  | luminance-matched | luminance-matched 1 |  |
| 5 | Luminance-matched receding side | centered at 45°, -9° (18° for LC6G2) | r/v = 10, 40, 70, 130, 310, 550 ms |  |  | luminance-matched 2 |  |
| 6 | Looming annulus side | centered at 45°, -9° (18° for LC6G2) | r/v = 10, 40, 70, 130, 310, 550 ms |  | looming annulus | looming annulus 1 |  |
| 6 | Looming annulus receding | centered at 45°, -9° (18° for LC6G2) | r/v = 10, 40, 70, 130, 310, 550 ms |  |  | looming annulus 2 |  |
| 7 | Bilateral Constant Looming Disc side | centered at 45°, 18° (for LC6G2) | 10,20,50°/s |  |  |  | ipsi-contra expanding |
| 8 | Small object motion stimulus progressive | azimuth 0° to 108°, -31.5°(31.5° for LC6G2) | 10,20°/s |  | small object | small object 1 |  |
| 8 | Small object motion stimulus regressive | azimuth 108° to 0°, -31.5°(31.5° for LC6G2) | 10,20°/s |  |  | small object 2 |  |
| 9 | Vertical bar motion stimulus progressive | azimuth 0° to 108° | 10,20°/s |  | vertical bar | vertical bar 1 |  |
| 9 | Vertical bar motion stimulus regressive | azimuth 108° to 0° | 10,20°/s |  |  | vertical bar 2 |  |

If visual-spatial information is wholly discarded in the LC6 glomerulus, then the LC6G neurons would be expected to have flat receptive fields that broadly cover the entire field of view of each eye. In contrast, we found that both LC6G1 and LC6G2 neurons responded to looming stimuli within a restricted portion of the visual field. The bilateral LC6G1 neurons responded with significant calcium increases to looming stimuli across most of the stimulated ipsilateral hemifield (Figure 3B, C). The strongest responses were to looming stimuli along the midline and above the equator (Figure 3B, C, and 3—figure supplement 1C; contours).

By comparison ipsilateral LC6G2 neurons did not respond to looming stimuli near the midline, or much below the equator (Figure 3B, C). Individual LC6G2 (Figure 3—figure supplement 1C) recordings also show responses that are more focal, restricted to a smaller receptive field than the LC6G1 responses. The LC6G2 neurons exhibited a spatially focused receptive field when imaged as a population, despite being a group of 4-6 neurons, indicating that the cells either share their spatial selectivity or are even more spatially selective individually. The differences between the LC6G downstream neuron receptive fields (Figure 3C), suggest that spatially selective read out of the LC6 neurons is implemented in the glomerulus.

### LC6 downstream neurons differentially encode visual stimuli

While LC6G1s and LC6G2s are functionally downstream of LC6, they responded differently to visual looming stimuli based on the region of the visual field in which they are presented. Given their strikingly different morphologies, we investigated whether these neurons differentially process visual information from LC6s. We presented a panel of visual stimuli (Table 2) to compare calcium responses of LC6G1s and LC6G2s to those of LC6. As previously reported, LC6 preferentially responded to dark looming stimuli, and also responded to the individual features of a looming stimulus (darkening in the luminance-matched stimulus and the edge motion in the looming annulus stimulus), but not to a receding dark stimulus or a bright looming stimulus (Wu et al., 2016). LC6 neurons also responded to the motion of non-looming objects (Figure 4B; response time series shown for slowest speed, tuning curves on the right for other speeds). The bilaterally projecting LC6G1 neurons responded to the looming related stimuli as well as the bar and small object motion stimuli in a manner that was very similar to the LC6 population (Figure 4B, C). This similarity of LC6 and LC6G1 responses extended to all stimuli presented, which is well illustrated by the scatter plot comparing the responses of the two cell types (Figure 4C, left). The regression line relating (normalized) LC6 to LC6G1 responses is close to the unity line (slope=1.05, Pearson’s correlation r=0.87), suggesting that LC6G1 may serve as a ‘summary’ of the LC6 populations’ visual responses across all stimuli. By contrast, the ipsilaterally projecting LC6G2s were much more selective for looming stimuli, responding to dark looming and to the edge motion of the looming annulus, but not to the other stimuli (Figure 4B). The dissimilarity of responses can also be observed from the comparison scatter plot (Figure 4C, right), where the points are more dispersed away from the unity line, and the linear regression shows a weaker correlation (slope=0.51, r=0.40). While LC6G1 relays the LC6s’ visual responses (presumably to the contralateral LC6 glomerulus) the LC6G2 neurons is more selective for looming than non-looming stimuli.

**Figure 4:**
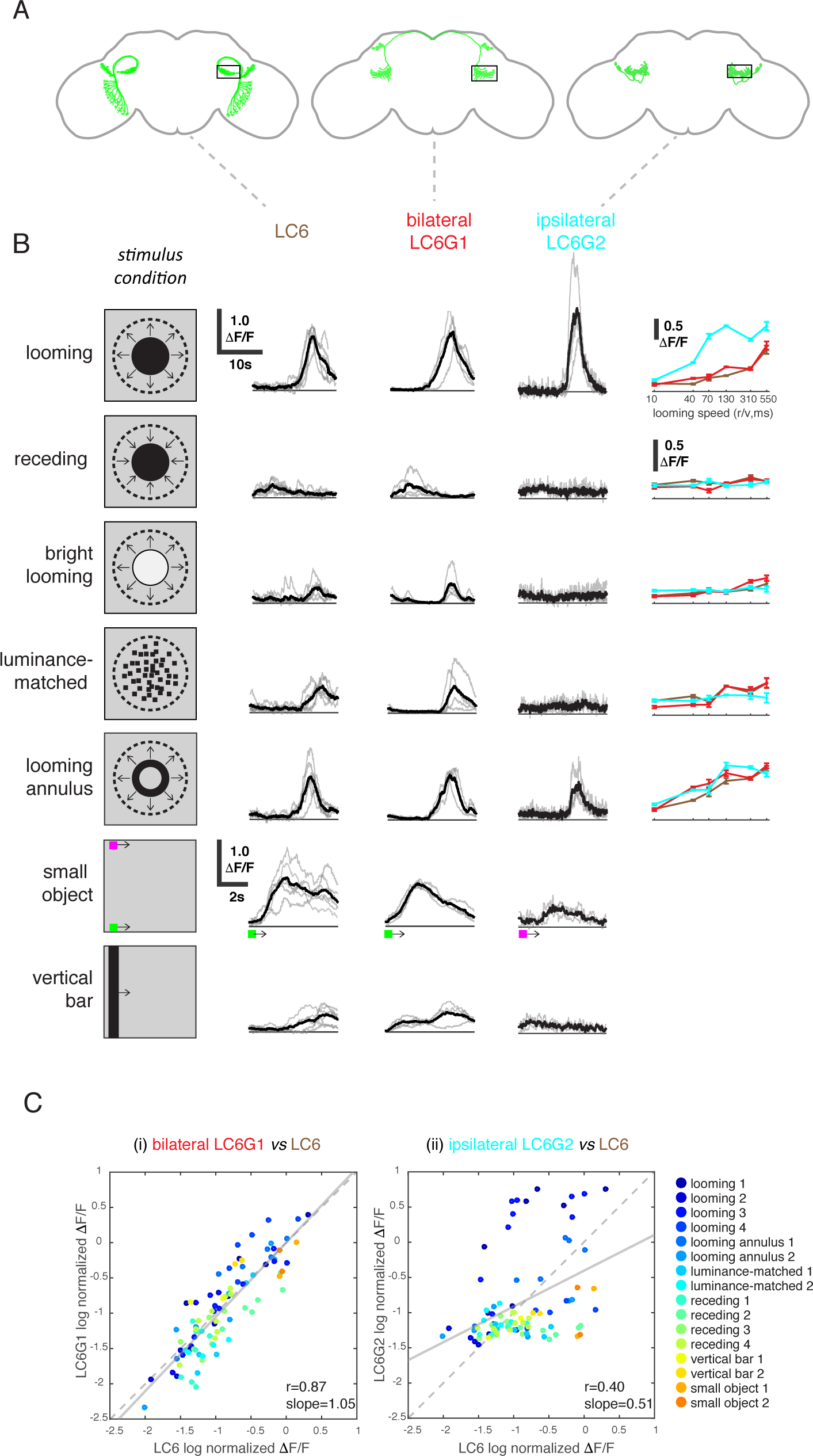
LC6 downstream neurons differentially encode visual stimuli. **(A)** Schematic of LC6, bilateral (LC6G1), and ipsilateral (LC6G2) projecting downstream cell types. Black rectangles indicate the ROI selected for GCaMP imaging; visual stimuli were presented to the ipsilateral eye. **(B)** Evoked calcium responses in LC6 and downstream neurons by visual stimulus in “stimulus condition” column. Times series are shown for the lowest speed (largest r/v time constant) in each stimulus category. Individual fly responses are in gray, and mean responses in black (LC6 N=6, LC6G1 N=4, LC6G2 N=3). The tuning curves show peak responses for each speed of the looming and looming-related stimuli (mean ± SEM). **(C)** Comparison between LC6 and the LC6G visual responses. LC6 and LC6G peak responses are log normalized and plotted against each other. LC6G1 (bilaterally projecting downstream) responses correlate well with that of LC6 (Pearson’s correlation, slope=1.05, r = 0.87, p=5.7×10^-29^), while LC6G2 (ipsilaterally projecting downstream) are not as well correlated (slope=0.51, r = 0.40, p=7.7×10^-5^). Note that most of the looming responses are above the unity line, while most of the non-looming responses are below it, demonstrating enhanced selectivity for looming stimuli. Linear fit is in black line, while X=Y (unity) line is in dashed gray. The data points are color coded by visual stimulus condition (detailed in methods). See Table 1 for detailed summary of fly lines used in this study and Table 2 for summary of all visual stimuli presented.

### LC6 neurons exhibit biased connectivity in the glomerulus

Our functional analysis establishes that LC6G neurons encode the visual-spatial information of looming stimuli from their inputs in the glomerulus, but anatomical analysis of the LC6 glomerulus from light microscopy images does not account for this selectivity (Figure 1). One parsimonious explanation is that this spatially selective readout of LC6 neurons could result from patterned subsets of synaptic connections between LC6 neurons and their downstream targets. To test this hypothesis, we mapped the LC6 neurons in a recently completed serial section transmission electron microscopy volume of the full adult brain (Zheng et al., 2018). From a single side (by convention, the right hand side, RHS) of the brain, we found 65 LC6 neurons (single example in Figure 5A), which we confirmed based on their morphology in the lobula, the distinctive axonal loop tract (shared with LC9), and the orientation of the axons in the glomerulus (Figure 5B). We then reconstructed all LC6 neurons, completing their axon terminals in the glomerulus, and tracing the major, but not the finest neurites in the lobula. Tracing the major dendritic branches in the lobula enabled us to computationally estimate the corresponding region of the visual field for each LC6 neuron (Figure 5C and methods). The coverage of the lobula is nearly uniform, but with a higher density toward the visual midline (Figure 5—figure supplement 1A). Using this mapping procedure, we constructed an ‘anatomical receptive field’ of LC6s, an RF estimate based only on the dendritic location and extent of each of the 65 reconstructed LC6 neurons (Figure 5D).

**Figure 5:**
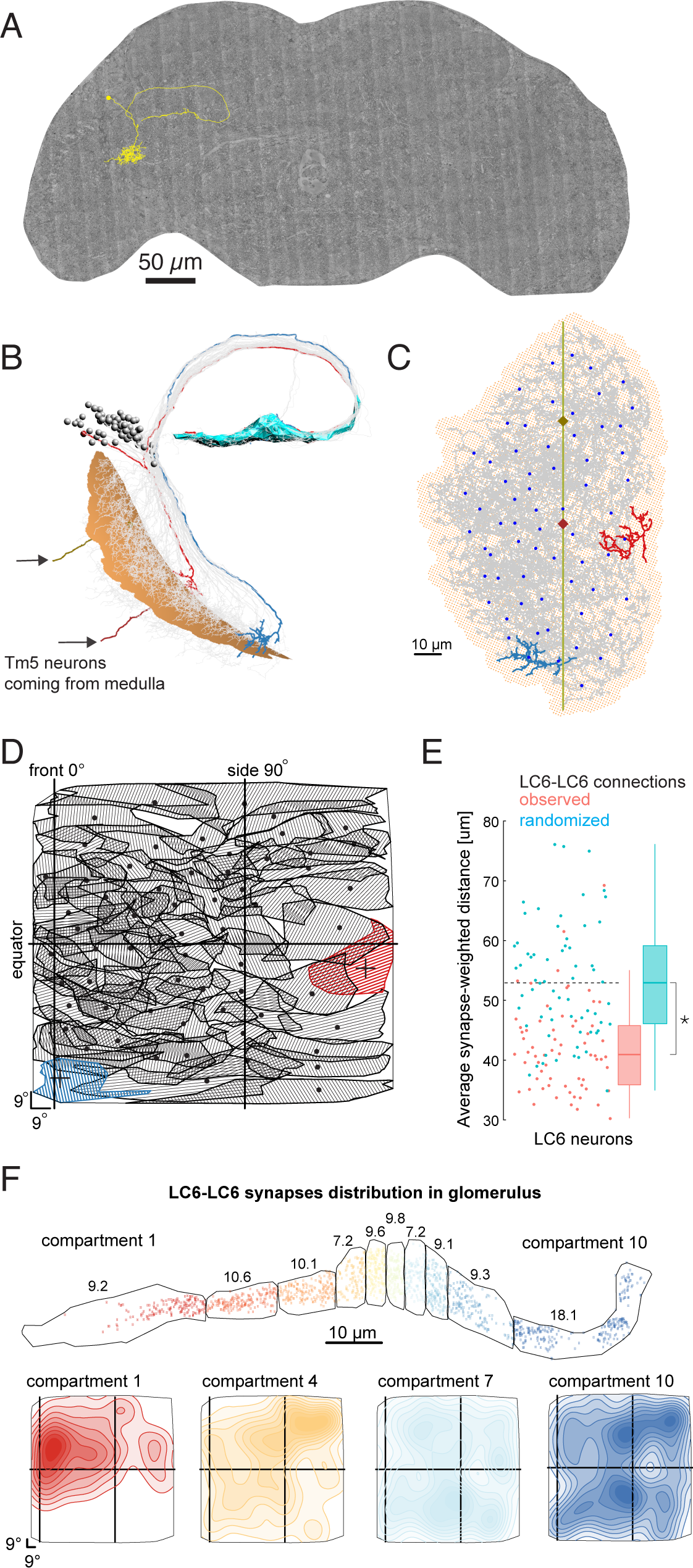
LC6 neurons exhibit biased connectivity in the glomerulus. **(A)** An example LC6 neuron found in Serial Section Transmission Electron Microscopy volume of the complete female brain. **(B)** All LC6 neurons on one side of the brain were traced into the glomerulus. Two example cells are highlighted in red and blue (same cells in **C** and **D**). All cells have dendrites mainly in lobula layer 4 (orange), and project to the LC6 glomerulus (cyan). Cell bodies are rendered as grey spheres. The projections of two Tm5 neurons are shown coming from the medulla side of the lobula. They were selected to identify the center (brown Tm5) and central meridian (both Tm5s) of the eye in **C**. **(C)** Projection view of the lobula layer. Traced LC6 dendrites (grey) are projected onto a surface fit through all dendrites (light orange). Blue disks represent centers-of-mass of each LC6 dendrites. The vertical line is the estimated central meridian, the line that partitions the eye between anterior and posterior halves, mapped onto the eye coordinate in **D. (D)** Estimate of dendritic field coverage of visual space (anatomical receptive field) for all LC6 neurons. Polygons are the estimated visual fields, and the boundary indicates the estimated boundary of the lobula (layer 4). The red example cell RF corresponds to posterior-viewing parts of the eye while the blue example cell RF corresponds to fronto-ventral viewing direction. **(E)** Analysis of LC6-LC6 connections in the glomerulus. LC6 neurons contact their dendritic neighbors with significantly greater probability than chance (shuffled for same number of synapses, but with visual-spatial relationships scrambled; Mann-Whitney U test, p = ∼ 0.015). The synapse-weighted distance is calculated by summing the products of distance and number of synapses between a given LC6 and the all others and the dividing by the total number of incoming synapses. **(F)** LC6-LC6 connections within 10 compartments of the glomerulus, selected to contain an equal number of LC6 pre-synapses. The numbers are the percentage of LC6-LC6 synapses within each compartment. Contour plot**s** show the spatial distribution of LC6-LC6 synapses in each compartment plotted in the coordinates of the field of view of one eye. The general trend is for middle compartments to exhibit broader synapse distribution, while those on either end of the glomerulus show more spatially restricted synapses. All compartments shown in Figure 5—figure supplement 1B.

We found and labeled many synapses between LC6 neurons in the glomerulus (1902 total, a mean of ∼29 synapses to each LC6 neuron from all other LC6 neurons, Table 3). We used this connectivity to look for a correlate of retinotopy in the glomerulus: do LC6 neurons preferentially synapse onto LC6 neurons with nearby dendrites (and thus spatially closer receptive fields)? When the distance between RF centers (measured across the lobula) was weighted by the number of synapses between the LC6 cells, we found that the average distance between connected neurons is significantly smaller than the distances after random shuffling of these connections (Figure 5E). Therefore, LC6 neurons bias their connectivity, such that they connect to their lobula neighbors with a distribution significantly smaller than produced by random connections.

**Table 3:**
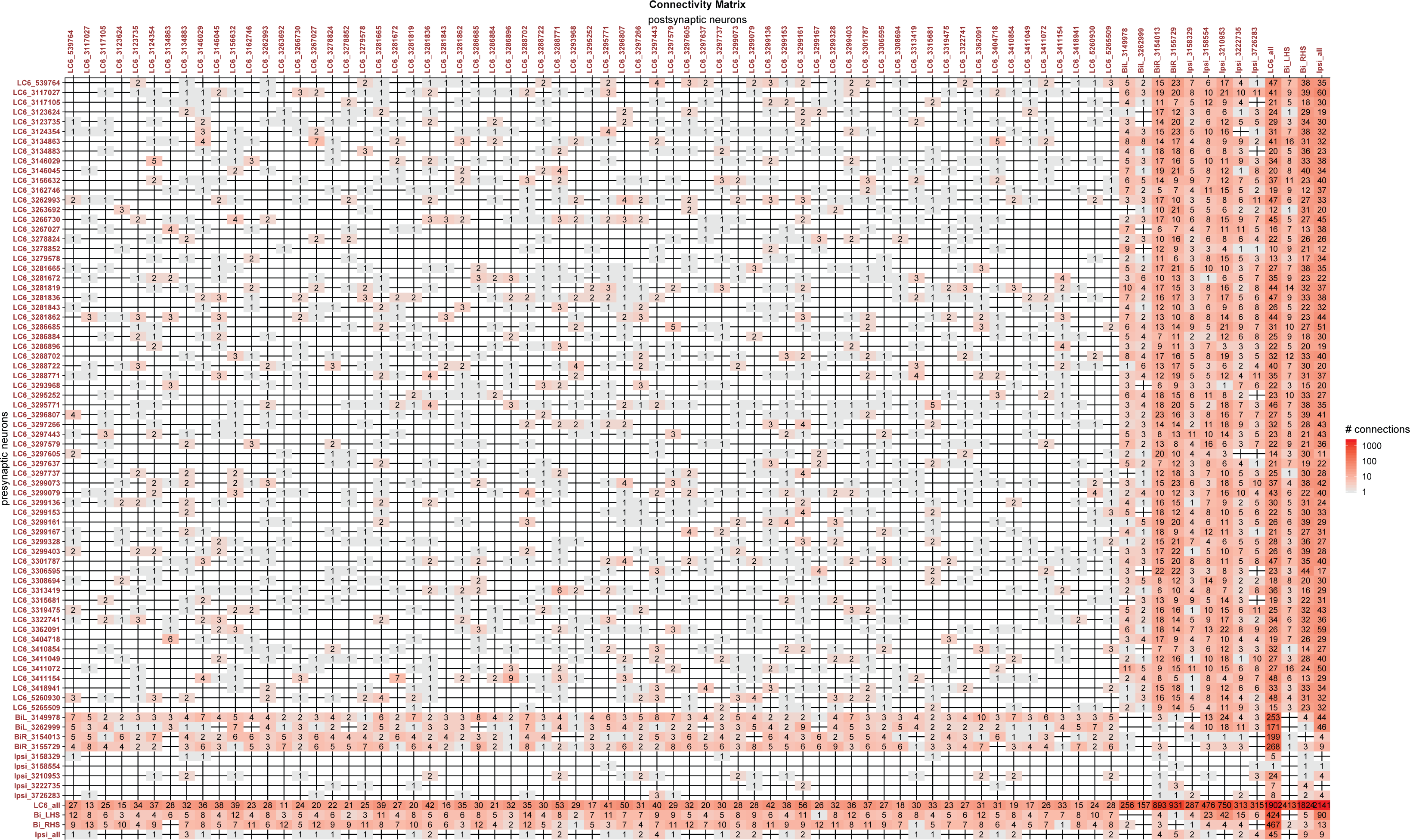
Connectivity matrix of LC6 and target neurons from EM data, related to Figure 6.

Since LC6 axons preferentially synapse onto other LC6 cells with neighboring dendrites, we next examined whether these synapses exhibit any spatial organization. To visualize this, we divided the glomerulus into 10 compartments along its long axis, each containing the same number of LC6 pre-synapses (across all postsynaptic cells; see ‘computational analysis of EM reconstruction’ in methods; Figure 5F). We then examined the spatial distribution of the LC6 input within each compartment as a contour plot in visual coordinates (see methods).

We found that the most lateral glomerulus compartment (#1) is biased towards representing the anterior visual field and the most medial glomerulus compartment (#10) is biased towards the posterior visual field. Most remaining compartments sample visual inputs more broadly, with peak responses close to the center of the eye (consistent with broad inputs covering much of the visual field; Figure 5—figure supplement 1B). This result was somewhat surprising—even though we could not find any obvious retinotopy of the LC6 axons in the glomerulus with light level anatomical analysis (Figure 1), the preference for synapsing onto LC6 neurons whose dendrites are neighbors in the lobula, as well as the spatial organization of these synapses in the glomerulus demonstrates that some retinotopy is maintained, but it is only observable in the organization of synapses.

### Connectivity-based readout of visual information in the LC6 glomerulus corroborates spatially selective responses of target neurons

The biased distribution of LC6-LC6 synapses demonstrates that some retinotopic organization exists in the glomerulus, but can the readout of LC6 inputs account for the spatially selective responses of the LC6G neurons? We manually reconstructed neurons within the LC6 glomerulus (by following, at random, neurites postsynaptic to LC6s), until we could match reconstructed neurons to the bilateral LC6G1 and ipsilateral LC6G2 cell types that were examined in Figure 3 and 4. We performed our analysis on the processes of four bilateral LC6G1 neurons (two each, per side of the brain) and on the unilateral axonal arbors of five LC6G2 neurons. These nine neuronal arbors were fully reconstructed and independently proofread (in the glomerulus; examples in Figure 6A). The connectivity between these nine arbors and the 65 LC6 neurons is summarized in Figure 6B (and detailed in Table 3). As expected from the morphology of these LC6 target neurons, and the functional connectivity (Figure 2), we found many synapses between the LC6 neurons and the right hand side (RHS) LC6G1 and LC6G2 neurons (mean number of synapses: ∼912 and ∼428, respectively). Perhaps more surprising is that we also found LC6 synapses onto the axons of the left hand side (LHS) LC6G1 neurons (mean of ∼207; LC6G1 (L) in Figure 6B); the neurons conveying contralateral visual inputs therefore also receive ipsilateral visual inputs. Further we find a similar number (∼223) of feedback synapses from the bilateral neurons of both sides onto LC6 cells.

**Figure 6:**
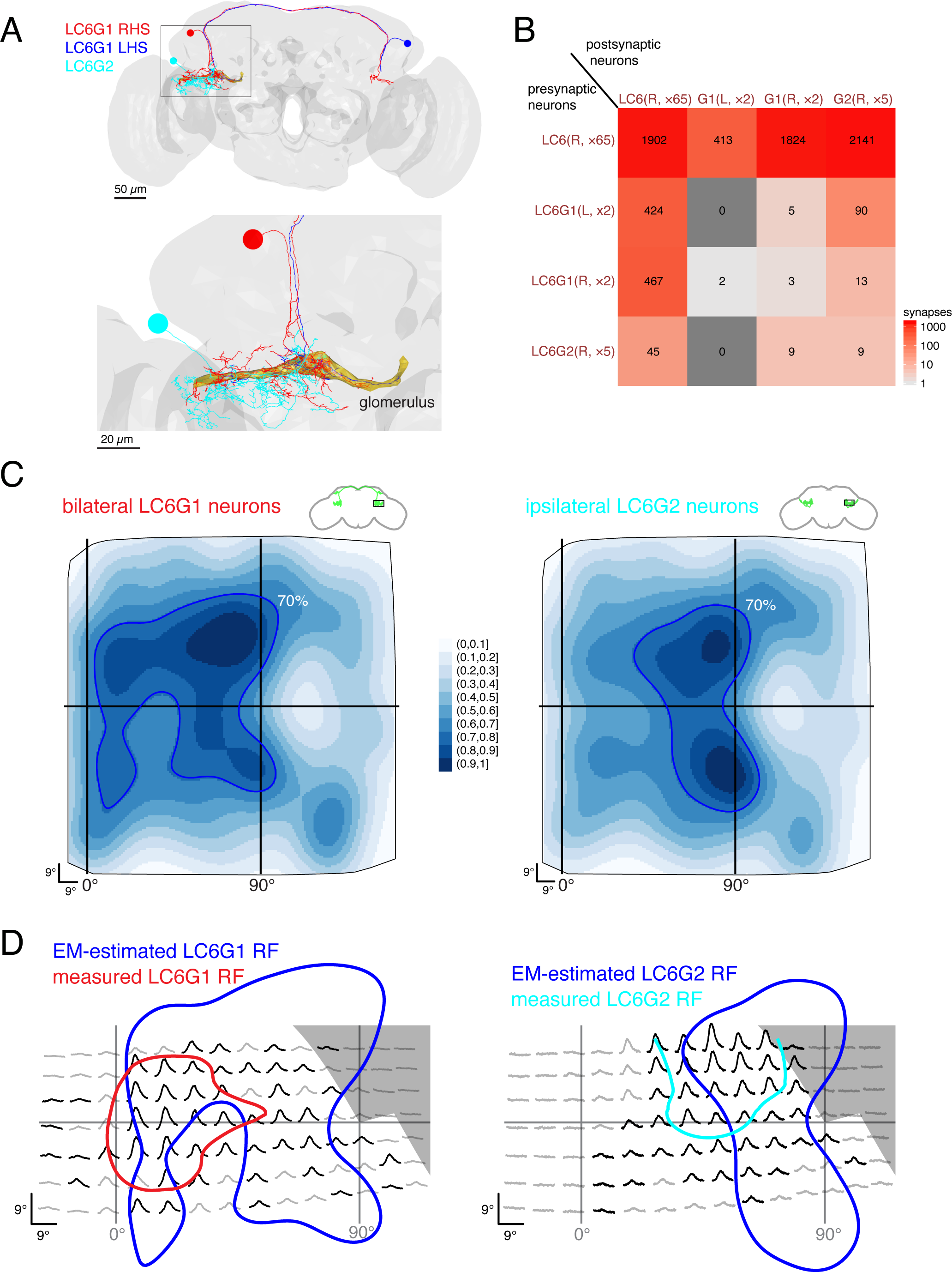
Connectivity-based readout of visual information in the LC6 glomerulus corroborates spatially selective responses of target neurons. **(A)** Bilaterally projecting LC6G1 and ipsilaterally projecting LC6G2 neurons were identified and reconstructed in the full brain EM volume. Individual neurons shown in Figure 6—figure supplement 1. **(B)** Summary connectivity matrix of LC6, LC6G1, and LC6G2 neurons, each cell type is grouped, with the side of the brain (R or L) and the number of neurons in each group labeled. See Table 3 for the complete connectivity matrix. **(C)** Estimated anatomical RFs of each target neuron type. Each LC6 neuron’s RF is scaled by the number of synapses to each individual target neuron, and then summed across all neurons of the same type (see methods for details). RFs for each individual target neuron shown in Figure 6—figure supplement 1. The 70% contour is highlighted with a dark blue line. **(D)** Anatomical RF (blue contour) overlaid onto functional RF measured *in vivo* (replotted from Figure 3C, but with the average 60% of peak contour shown here). Further comparisons between these cell types and between EM and functional estimates of the RFs are in Figure 6—figure supplement 2.

With the synapses between LC6 neurons and their targets mapped, we could finally attempt to correlate the connectivity-based readout of visual information with our functional measurements of spatial receptive fields (Figure 3). Using gaussian approximations for the LC6 anatomical RFs (Figure 5D), we estimated the anatomical RFs of the individual target neurons by summing over all input LC6 RFs, scaled by the number of synapses. We summed the estimated RFs for each cell type across individual cells to compare with the population measurements during calcium imaging experiments (Figure 6—figure supplement 1, see methods). These summed estimated RFs are shown for LC6G1 and LC6G2 (Figure 6C) across the field of view of the entire right eye. This analysis shows several features that agree with our functional RF mapping of these cell types (Figure 3). The LC6G1 receptive field appears broader and has large responses close to the frontal midline, while the LC6G2 neurons have their strongest responses in a smaller zone, away from the midline. To generate a more direct comparison, we aligned the functional RFs measured through *in vivo* calcium imaging (generated with stimuli that partially covered the visual field; Figure 3C) to the 70% contour of the anatomical estimate for each reconstructed cell type (Figure 6D). Considering that these estimates of the RF of each cell type were generated with very different methods, and accounting for the approximations required to align the two data sets, we find this level of agreement between these RFs to be substantial. For example, the spatially broader responses of the LC6G1 cells closer to the frontal midline and the much narrower responses of the LC6G2 cells away from the midline (0° azimuth), are readily observable in both measures of the RF.

Could the RFs estimated for the LC6G neurons be explained by the localization of synapses within the glomerulus (and the visual fields they represent, Figure 5F)? Further analysis of where the LC6Gs collect their inputs within the glomerulus (Figure 6—figure supplement 2), shows that the establishment of the spatially selective RFs is not a trivial reflection of synapse location within the glomerulus, since the differences between LC6G1 and LC6G2 visual inputs are present throughout the glomerulus. Taken together this connectomic analysis of the LC6 glomerulus clearly establishes that spatially selective readout by optic glomerulus neurons is accomplished using biased connectivity within a structure in the central brain where the neuronal arbors themselves are not retinotopically arranged.

### Interconnections between the LC6 glomeruli support contralateral suppression of visual responses to looming stimuli

One prominent feature of the fly nervous system is that brain compartments are often connected—directly or indirectly—to their symmetric compartment across the hemispheres (Shih et al., 2015). What role does this flow of information play? Of the LC6G neurons we identified (Figure 1), only the LC6G1 neurons consist of a symmetric population of neurons that cross the midline and innervate both LC6 glomeruli. The set of neurons whose interconnections we completely reconstructed within the LC6 glomerulus are fully detailed in Table 3 and summarized in Figure 7A. The LC6G1 neurons from both hemispheres are both pre- and post-synaptic in the LC6 glomerulus. The RHS LC6G1 neurons receive ∼4.4 times more synapses from LC6 neurons in the RHS glomerulus than the LHS LC6G1 neurons, however, these neurons from both sides of the brain provide a similar number of synapses back onto LC6 neurons. Only the contralateral (LHS) LC6G1 neurons synapses onto the ipsilateral LC6G2 neurons, while the LC6G2 neurons do not synapse onto either of the LC6G1 neurons.

**Figure 7:**
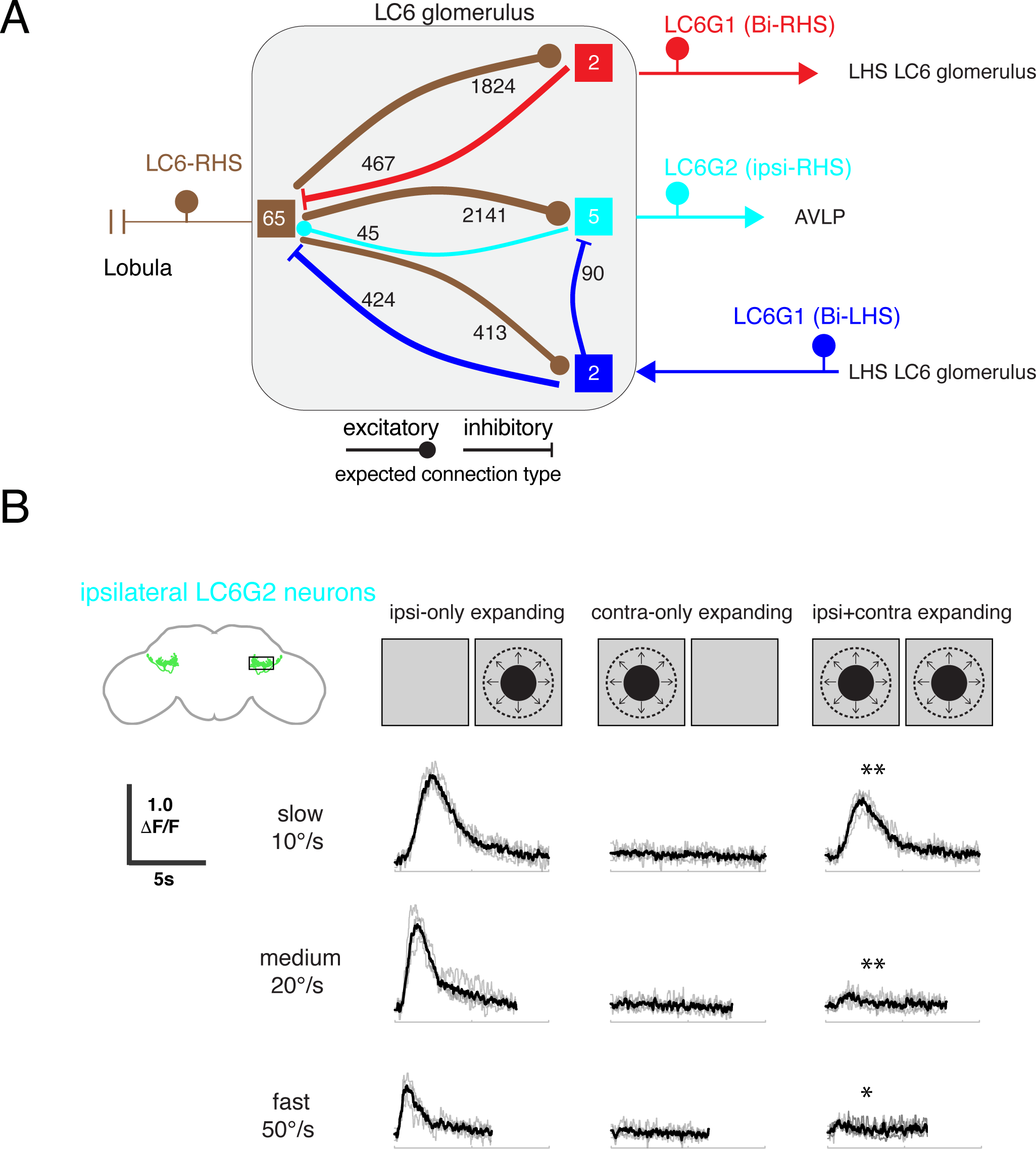
Interconnections between the LC6 glomeruli support contralateral suppression of visual responses to looming stimuli. **(A)** A proposed microcircuit between LC6s, LC6G1, and LC6G2. Synapse counts are shown for observed connections in the EM reconstruction (above threshold of 15 synapses). Each colored box indicates one of the 4 groups of neurons in the connectivity matrix (Figure 6B), and the numbers of cells within each group are listed within each box. Based on expression of neurotransmitter markers (Figure 1—figure supplement 3), LC6 and LC6G2 are expected to be excitatory (indicated by the ball-shaped terminal), and LC6G1s are expected to be inhibitory (bar-shaped terminal). Within the glomerulus the line thickness indicates the number of synapses in each connection type (logarithmically weighted), which is also noted next to each connection. Outside of the glomerulus, arrowheads are used to depict the direction of the projection, away or towards the glomerulus. **(B)** Evoked calcium responses in LC6G2 by variations of bilateral looming stimuli. Looming stimuli are centered at ±45°. Responses for three different, constant speeds of looming. Mean responses from individual flies are in gray and mean responses across flies in black (N=4). The LC6G2 response is significantly reduced by the simultaneous presentation of a contralateral looming stimulus at all speeds (p = 2.2×10^-3^, p = 4.8×10^-3^, p = 2.3×10^-2^ at 10, 20, 50°/s respectively, two-sided paired t-test). See Table 2 for summary of all visual stimuli presented.

By combining the connectivity of these three cell types together with the neurotransmitter expression profiles (Figure 1—figure supplement 3), a simple microcircuit emerges (Figure 7A). The flow of information from the contralateral LC6 glomerulus via the putatively inhibitory LC6G1 neurons may serve a gain control function within the glomerulus using both ipsilateral and contralateral responses of LC6 neurons to dampen strong aggregate responses from the inputs to the glomerulus. Furthermore, the contralateral (LHS) LC6G1 neurons directly synapse onto the ipsilateral (RHS) LC6G2 neurons, potentially suppressing their stimulus responses as a consequence of contralateral visual inputs.

To test this proposal, we recorded the calcium responses of the ipsilateral LC6G2 neurons while presenting looming stimuli to each eye (Figure 7B). As expected from the results in Figure 4, when a single looming stimulus was shown, LC6G2 neurons responded to the ipsilateral, but not the contralateral, presentation (Figure 7B, columns 1,2). When both sides were simultaneously stimulated with looming, the responses were significantly reduced at all tested speeds (as compared to the ipsilateral only stimulus condition; Figure 7B, column 3). This result agrees well with our working model in which the responses of the LC6G2 glomerulus projection neurons are suppressed by the presence of the preferred (looming) stimulus from the contralateral eye, mediated by inhibition from the bilaterally projecting LC6G1 neurons. Together these results suggest that contralateral visual inputs further enhance the spatial selectivity of these LC6 projection neurons.

## Discussion

In this study we investigated the circuitry downstream of the looming responsive LC6 visual projection neurons. We used light-level anatomical analysis to identify candidate downstream neurons (Figure 1 and 1—figure supplement 1) and then used intersectional genetic methods to establish driver lines that precisely target each of these putative downstream target neurons (Figure 1—figure supplement 2). We used these driver lines to show that five of these cell types are functionally connected to LC6 neurons (Figure 2). We then selected two of these cell types, the bilaterally projecting LC6G1 neurons and the ipsilateral LC6G2 projection neurons for further functional studies. We found that these two cell types differentially encode looming stimuli (Figure 3 and 4). The LC6G1 neurons responded to looming stimuli over a large part of the visual field and showed nearly identical stimulus selectivity as the LC6 neurons. By contrast, the LC6G2 neurons responded to looming stimuli within a more restricted region of the visual field, and showed enhanced stimulus selectivity, preferentially encoding looming stimuli, while being less responsive to non-looming stimuli than their LC6 inputs.

The organization of visual projection neuron axons into glomeruli in the fly brain has been the subject of significant speculation. One popular proposal, inspired by the loss of retinotopy within the glomeruli, is that these structures represent a transformation from a visual, spatial ‘where’ signal to a more abstract ‘what’ representation (Mu et al., 2012; Strausfeld et al., 2007; Wu et al., 2016). However, we observed that the LC6 glomerulus target neurons are spatially selective (Figure 3), and so we pursued a connectomic reconstruction of the LC6 glomerulus and showed that while some retinotopic information is present in the LC6 glomerulus (Figure 5), the spatial receptive fields of the LC6G neurons are established by biased synaptic connectivity (Figure 6). Finally, our connectomics analysis clarified the flow of information between these target neurons and LC6 (Figure 7).

From this functional and anatomical evaluation of the circuit, the picture that emerges is of glutamatergic (putative inhibitory) LC6G1 neurons that serve as interneurons within and between the glomeruli, reporting the summed LC6 activity, which is used to suppresses high (or broad) levels of activation—as would be encountered during forward locomotion in a cluttered environment. In contrast, the cholinergic (excitatory) LC6G2 projection neurons, convey the detection of looming stimuli to the AVLP, a deeper brain area. These neurons show enhanced selectivity for looming stimuli within one eye (Figure 4), but also receive inhibitory inputs (at least in part mediated via LC6G1 neurons, Figure 7) that would further serve to preserve responses to looming stimuli while suppressing responses to global visual stimulation.

### Organization of visual space in the LC6 glomerulus

We have investigated how visual-spatial information, representing different retinotopic coordinates, is organized in the LC6 glomerulus through light and EM anatomy. From confocal images, the LC6 axons (Figure 1C’) do not show any retinotopic organization along either the long or short axes of the glomerulus. However, from our EM reconstruction of the LC6 glomerulus (Figure 5 and 6), we found at least three manifestations of spatial organization. (1) We found that neighboring LC6s in the lobula preferentially form axo-axonic connections in the glomerulus (Figure 5E). (2) The location of these LC6-LC6 synapses were found to have a bias: synapses from LC6s with dendrites corresponding to the anterior visual field tend to cluster at the distal end of the LC6 glomerulus, while the synapses from LC6s with dendrites that correspond to posterior eye regions, tend to cluster at the proximal end (Figure 5F). (3) The LC6 downstream neurons receive selective connections from LC6 neurons in the glomerulus. The synapses of LC6G1 and G2 were distributed broadly across the LC6 glomerulus (Figure 6—figure supplement 2), however, by forming higher numbers of synapses onto subsets of the LC6 neurons throughout the glomerulus, these target neurons access inputs corresponding to restricted regions of the visual field (Figure 6 and 6—figure supplement 1). Taken together, we find only weak evidence for a continuous spatial organization (“retinotopy”) in the LC6 glomerulus at the light or the EM level (Figure 5). Instead we find that the largely overlapping processes of individual LC6G neurons, through selective connectivity, are able to integrate visual-spatial information from a restricted region of the eye. This situation is somewhat analogous to the well-studied T4 and T5 neurons, where the dendrites of these neurons, within retinotopically organized neuropils, prominently overlap with many cells of the same and different subtypes, and yet establish precise connections to specific cell types only at specific dendritic locations (Shinomiya et al., 2019). While visual-spatial readout is supported by connectivity in the glomerulus, we find strong evidence for a coarsening of the visual representation, as neurons exhibit increasingly larger field of views, with all target neurons integrating from the majority of LC6 neurons. Further work will be required to determine if these transformations described in the LC6 glomerulus are a general circuit strategy employed within the other optic glomeruli.

What benefits, if any, might this peculiar glomerulus structure serve for the organization of visual projection neurons? One prevailing view is that retinotopy affords wiring efficiency for downstream cells, provided they benefit from reading out information from neighboring visual regions (Chklovskii and Koulakov, 2004). On the other extreme, if the readout strategy requires non-spatial, random access to the entire visual field, then a compact, ball-like structure would minimize wire length. But if the purpose of some downstream neurons is to access all of the inputs (e.g. LC6G1) while others integrate from only a small neighborhood of the visual field (e.g. LC6G2), then the organization we observe in the LC6 glomerulus presents a sensible compromise—a compact structure which minimizes overall wire length for ‘global integrating’ neurons, while still allowing retinotopic readout, albeit somewhat coarsened, to support spatial vision.

### Spatial detection of visual looming

Given that LC6 downstream cells respond to looming stimuli (Figure 4), how might the organization of visual-spatial information in the LC6 glomerulus contribute to the fly’s ability to detect looming features? At the level of LC6 population, the neighboring LC6-LC6 connections might contribute to the detection of looming. We only described axo-axonic connections, but dendro-dendritic connections between overlapping neurons are also expected. Lateral facilitation should enhance responses to looming stimuli by this population of non-directionally selective cells. The first cells that are excited by a small dark object would facilitate the neighboring cells, and as the object grows in size, the surround facilitation would amplify responses while spreading as an outwardly radiating wave. At the same time, this scenario would not enhance responses to a receding object, which is consistent with the LC6 population responses (Figure 3, (Wu et al., 2016)). At the level of LC6 downstream cells, we have shown that LC6G1 and LC6G2 display different levels of selectivity for looming features (Figure 4). While determining the mechanism of this enhanced selectivity for looming is beyond the scope of this paper, the observation of this selectivity enhancement strongly suggests that the LC6 pathway is specialized for the detection of looming stimuli, since targets of this pathways are even more selective for looming stimuli than the visual projection neurons.

Since the LC6G1 and LC6G2 cells exhibit different receptive fields (Figure 3), it is likely that the downstream cell types act as different filters for particular looming features in particular regions of space. The kind of spatial and stimulus selectivity we measured for LC6G2 neurons (Figure 3 and 4) could support directional escapes, where approaching stimuli elicit escape take-offs in roughly the opposite direction (Card and Dickinson, 2008). Further, it seems essential for the fly to discriminate between the ground looming upwards in the ventral visual field and an approaching object in the medial visual field. These events would require triggering significantly different motor programs, such as landing or evasion. Therefore, the organization we observe in LC6 target neurons (especially LC6G2) might provide an efficient circuit logic to transform a purely sensory input into a signal that drives specific motor reactions, such as directed escape. From this perspective, the coarsened visual-spatial readout in the LC6 glomeruli may be the expected feature of an intermediate step in a sensory-to-motor transformation.

### The role of contralateral suppression

Through our EM reconstruction, we found interconnectivity between LC6, LC6G1, and LC6G2 within the glomerulus. Together with their neurotransmitter identities (Figure 1— figure supplement 3), we proposed a simple circuit within the LC6 glomerulus capable of contralateral suppression (Figure 7). We tested this hypothesis using bilateral looming stimuli while imaging from LC6G2 and found that the stimulus-evoked responses were consistent with the proposed circuit mechanism. Contralateral suppression was already implied from light level anatomy that showed the bilateral downstream cell type specifically contacted the same glomeruli in both hemispheres. This type of connection between glomeruli is common; based on our analysis of Janelia’s GAL4 collection, we expect that most if not all optic glomeruli are connected by one or more bilaterally projecting interneuron types. Also, contralateral suppression has been observed in other contexts and brain regions in *Drosophila*, such as detection of visual bilateral bar stimuli (Sun et al., 2017) or wind direction (Suver et al., 2019), in the latter case being implemented by bilaterally projecting cell types of similar morphology to LC6G2. This suggests that the use of bilaterally projecting cell types might be a common strategy used for contralateral suppression in the *Drosophila* CNS. In addition to being common, it is also sensible (and is related to the engineering principle of designing amplifiers with ‘common mode rejection’). A major goal of sensory systems appears to be the detection and localization of specific sensory cues, and yet in the natural world cues are rarely discrete, they emerge in complex mixtures across space and time. From this perspective, one of the simplest methods to identify a specific, localizable threat, such as an approaching object, is to compare whether this detection event is restricted to one side of the animal, or whether it is prominent on both sides. We propose that contralateral suppression is likely to be a basic strategy used in establishing stimulus selectivity.

### Outlook

Here we described three types of spatial organization within the LC6 glomerulus synthesizing data from *in vivo* calcium imaging, functional connectivity, and light and EM level anatomy. Using cell type specific driver lines (Wu et al., 2016) and the recently completed full adult fly brain EM volume (Zheng et al., 2018), we correlated neurophysiological measurements with EM connectivity between identified cell types. As a result, we have described how visual-spatial information is read out of a neuropil that lacks retinotopic organization and proposed a circuit between LC6 and LC6 downstream cells that enables contralateral suppression. As EM-level connectomes become available, the approach we used of integrating functional analysis with circuit mapping should greatly accelerate the discovery of mechanisms for diverse circuit computations.

## Methods

### Fly lines: split-GAL4 generation and anatomical analyses (light microscopy level)

To construct split-GAL4 lines for potential LC6 target neurons we first selected candidate AD and DBD hemi drivers by visually searching images of GAL4 driver expression patterns (Dionne et al., 2018; Jenett et al., 2012b; Tirian and Dickson, 2017). Typically, we tested several candidate split-GAL4 combinations for each target cell type. The AD and DBD hemi drivers we used are from published collections (Dionne et al., 2018; Tirian and Dickson, 2017). A detailed description of split-GAL4 hemi drivers is also available online (https://bdsc.indiana.edu/stocks/gal4/split_intro.html) and the cell-type specific split-GAL4 lines generated in this study together with selected images of their anatomy will be made available at https://www.janelia.org/split-GAL4. All fly lines used throughout the manuscript are summarized in Table 1.

To visualize the overall expression patterns of the split-GAL4 driver lines, we used previously described reporter transgenes (pJFRC51-3XUAS-IVS-Syt::smHA in su(Hw)attP1 and pJFRC225-5XUAS-IVS-myr::smFLAG in VK00005 (Nern et al., 2015) and methods (Aso et al., 2014; Wu et al., 2016). Detailed protocols are also available online (https://www.janelia.org/project-team/flylight/protocols under “IHC - Anti-GFP”, “IHC - Polarity Sequential” and “DPX mounting”). For stochastic labeling of individual cells, we used standard protocols for Multicolor Flp-out (MCFO) (Nern et al., 2015). Briefly, split-GAL4 lines were crossed to MCFO-1 and adult progeny heat-shocked and immunolabeled with antibodies against FLAG, HA and V5 epitopes as described (Nern et al., 2015). Detailed protocols are also available online (https://www.janelia.org/project-team/flylight/protocols under “IHC - MCFO”. We examined expression of the neurotransmitter markers GAD1, VGlut and ChaT by Fluorescence in situ Hybridization (FISH) using published protocols and probe sets (Meissner et al., 2019). We note that in general we used the same driver lines across out experiments, with one noteworthy exception. We developed a more specific driver line for LC6G5 (SS25111) in the later stages of this project. This line was used only for the FISH experiments (Figure 1—figure supplement 3, Table 1), where interpretation of results benefits from the cleanest available driver line.

Images were acquired on Zeiss LSM 710 or 800 confocal microscopes with 20x 0.8 NA or 63x 1.4 NA objectives. We used Fiji (http://fiji.sc) to generate maximum intensity projections of driver patterns and sections of FISH images. For these images, adjustments were limited to changes of brightness and contrast. Most other anatomy panels show composites of multiple registered images, which were generated using a recently described template brain (Bogovic et al., 2018). To more clearly display the cells of interest images used in these panels were, in some cases, manually edited to exclude additional cells or background present in the original images. Editing and assembly of these composites was mainly done using FluoRender (http://www.sci.utah.edu/software/fluorender.html).

### Fly lines: functional connectivity

For the functional connectivity experiments (Figure 2), in order to express Chrimson (Klapoetke et al., 2014) and GCaMP6f (T. Chen et al., 2013) in different neurons, LexA/LexAop and split-GAL4/UAS systems were simultaneously used in the same animal. For exploring specific downstream candidate neurons, a LexA line containing the LC6 pattern (R42E06 in JK73A, (Knapp et al., 2015)) was used to drive Chrimson in LC6, while the split-GAL4 lines (ss0825, ss2036, ss2099, ss3690, ss2409, ss3641) were used to drive expression of opGCaMP6f in candidate LC6 downstream neurons (summarized in Table 1). Flies were reared under standard conditions (60% humidity, 25°C) on a cornmeal agar diet supplemented with retinal (0.2mM) in vials that were wrapped in foil to keep flies in the dark to prevent spurious activation of Chrimson by ambient light. Flies were collected following eclosion and held under the same rearing conditions until experiments were performed.

### Fly lines: calcium imaging

Cell-type specific expression of the fluorescent calcium indicator GCaMP6m (T. Chen et al., 2013) was achieved using the split-GAL4/UAS expression system (Luan et al., 2006; Pfeiffer et al., 2010). The GAL4 driver lines were constructed using methods described above. All flies used for calcium imaging experiments were reared under standard conditions (25 °C, 60% humidity, 12 h light/12 h dark, standard cornmeal/molasses food), and all imaging experiments were performed on females 3-6 days post-eclosion. To image from LC6 and its downstream targets, split-GAL4 driver lines [LC6: OL0070B, LC6G1(bilaterally projecting downstream): ss0825, LC6G2(ipsilaterally projecting downstream): ss2036] were crossed to pJFRC7-20XUAS-IVS-GCaMP6m in VK00005 (DL background) effector line. In a subset of the recordings the LC6 neurons, and thus the glomerulus, were labeled with tdTomato (to aid in selecting appropriate imaging plane). The lines used in each experimental Figure panel are summarized in Table 1.

### Functional connectivity: preparation, experimental details, and data analysis

Brains from female adult flies 1-3 days post-eclosion, were isolated by dissecting the head in a saline bath (103mM NaCl, 3mM KCl, 2mM CaCl2, 4mM MgCl2, 26mM NaHCO3, 1mM NaH2PO4, 8mM trehalose, 10mM glucose, 5mM TES, bubbled with 95% O2 / 5% CO_2_). The brain was then placed on a poly-lysine coated coverslip (neuVitro, Vancouver, WA, GG-12-PDL) posterior side up and perfused with saline (same composition as above, 21°C) (Figure 3A). Images of the brain were acquired using a two-photon microscope (Custom made at Janelia by Dan Flickinger and colleagues, with a Nikon Apo LWD 25× NA1.1 water immersion objective #MRD77225). The standard imaging mode was a 512 × 512 image at 2.5 frames/s, and a ∼353 μm × ∼353 μm field of view (∼0.69 μm × ∼0.69 μm / pixel). The sample was imaged using a near-infrared laser (920nm, Spectra Physics, Insight DeepSee) that produced minimal collateral activation of Chrimson at our typical imaging power.

The light-gated ion channel Chrimson was activated by 590nm light (Thorlabs M590L3-C1) presented through the objective. Photoactivation light was delivered in a pulse train that consisted of six 1s pulses (within each 1 s pulse: square-wave modulation at 50 Hz, 10% duty cycle, 30s inter-pulse interval). Two stimulation protocols, in succession, were used on each brain, the first (Figure 2) in which the light intensity increased for each of the six pulses (0.12, 0.24, 0.48, 0.72, 0.96, 1.21 mW/mm^2^; measured using Thorlabs S170C microscope slide light power meter) and the second (Figure 2—figure supplement 1) in which the light intensity was kept constant for each of the six pulses (0.48mW). Stimulation light was spatially modulated using a DMD (Digital Micromirror Device, Texas Instruments, DLP LightCrafter v2.0), and was restricted to one or both LC6 glomeruli depending on the experiment (Figure 2: left glomerulus only stimulation and ROI on left glomerulus region; Figure 2—figure supplement 1 left and right glomerulus only stimulation and ROI on left glomerulus region). This spatial restriction limited the activation of other few non-specific cells labeled with the same driver line.

Image registration was conducted using code from the Thunder package (https://github.com/thunder-project/thunder (Freeman et al., 2014)).

The calcium responses of candidate LC6 downstream neurons to photoactivation were measured by calculating the ΔF/F for a manually drawn region of interest (ROI) in the imaging plane, which covered the largest section of its dendritic arborization. The ΔF/F was taken as (F-F0)/F0 where F is the instantaneous mean fluorescence of the ROI and F0 is the baseline fluorescence of the ROI. The baseline fluorescence was taken as the 10th percentile of the stimulation protocol period. Peak responses from each fly for both protocols were taken, and the mean ± SEM is shown as a stimulus response curve in Figure 2 and 2—figure supplement 1.

### *In vivo* 2-photon calcium imaging: preparation

The imaging preparation was nearly identical to what we have described in a recent publication (Strother et al., 2014). Briefly, flies (mostly female, but some male) were cold anesthetized and tethered to a fine wire at the thorax using UV-curing adhesive. The two most anterior legs (T1) were severed and glued down along with the proboscis to prevent grooming of the eyes and to immobilize the head. Tethered flies were glued by the head capsule into a fly holder and after addition of saline (103mM NaCl, 3mM KCl, 1.5mM CaCl2, 4mM MgCl2, 26mM NaHCO3, 1mM NaH2PO4, 8mM trehalose, 10mM glucose, 5mM TES, bubbled with 95% O2 / 5% CO2; final Osm=283, pH=7.3; modified from (Wilson and Laurent, 2005)) to the bath, the cuticle at the back of the head was dissected away to expose the brain. Muscles 1 and 16 (Demerec, 1965) were severed to reduce motion of the brain within the head capsule, and the post-ocular air sac on the imaged side of the brain was removed to expose the optic glomeruli. The right side of the brain was always imaged (unless otherwise stated).

### *In vivo* 2-photon calcium imaging: microscopy

The optic glomeruli were imaged using a two-photon microscope (Bruker/Prairie Ultima IV) with near-infrared excitation (930 nm, Coherent Chameleon Ultra II) and a 60x objective (Nikon CFI APO 60XW). The excitation power was 10-20 mW at the sample. The long excitation wavelength is not visible to the fly. Imaging parameters varied slightly between experiments but were within a small range around our typical acquisition parameters: 128 x 90-pixel resolution, and ∼10 Hz frame rate (10.0-10.5 Hz). LC cell axon calcium data were collected from single planes selected to capture a consistently large slice of each glomerulus. In several experiments the glomerulus was labeled with tdTomato and located with a separate red-channel detector. The filter sets used in the detection path were HQ525/70 bandpass (Chroma) and 510/42 bandpass (Semrock) for the green-channel, HQ607/45 bandpass (Chroma) and 650/54 bandpass (Semrock) for the red-channel.

### *In vivo* 2-photon calcium imaging: visual stimulation

Visual stimulation details closely follow the methods used in our previous study (Wu et al., 2016). Flies were placed near the center of a modular LED display (Reiser and Dickinson, 2008) on which visual stimuli were presented (Figure 3A). The display consists of 574 nm peak output LEDs (Betlux ultra-green 8×8 LED matrices, #BL-M12A881UG-XX) covered with a gel filter (LEE #135 Deep Golden Amber) to greatly reduce stimulus emission at wavelengths that overlap with those of GCaMP emission, resulting in emission range of 560-600nm (measured by Ocean Optics USB4000-UV-VIS spectrometer). The display was configured to cover 60% of a cylinder (32 mm × 32 mm panels. forming 12 sides of an extruded 20-sided polygon, approximating a cylinder, each column is 4 panels high; total resolution is 32 pixels × 96 pixels).

Since the head is fixed into the fly holder with an approximately constant orientation (relative to horizontal, Figure 3—figure supplement 1A), the display was tilted to roughly match this angle. The projection of the display onto the flies’ field of view was estimated (Figure 5B). First careful measurements (angles and distances) of the experimental setup were made, and then an optimization procedure was used to more precisely estimate the angle and position of the tethered fly within the cylindrical display. Using these measurements, the location of every pixel in the display was mapped to 3-dimensional cartesian coordinates relative to the fly position and orientation, and then converted to a pixel-map in spherical coordinates.

Additionally, the occlusion by the holder to which the fly is mounted was also transformed into spherical coordinates. The field of view of *Drosophila melanogaster* was estimated based on Andrew Straw’s digitized eye map (http://code.astraw.com/drosophila_eye_map/), which was in turn based on E. Buchner’s pseudo-pupil mapping reported in his Diplom (Buchner, 1971). The composite projection for the estimated positions of every pixel in the display, the position of every ommatidia, and the occlusions due to the holder are shown in Figure 3—figure supplement 1B. As the fly was positioned slightly closer to the front of the display from the center of the cylinder, the spacing between pixels is not constant, but averages to 2.5° along the eye equator. The slight offset between the fly’s frontal midline and the center of the display, as well as the small offset between the fly eye’s equator and the display’s horizon capture our best attempt at an accurate alignment. It is important to note that the fly holder occludes large portions of the fly’s dorsal and lateral field of view. We believe this alignment to be accurate to within 5°-10°, or the spacing between 1-2 ommatidia. Code for this transformation is posted at https://github.com/reiserlab/LC6downstream.

Looming stimuli were used to map the receptive fields of the LC6G neurons (stimulus type 1, Table 2). Dark discs expanding from ∼4.5° to ∼18°, at constant velocity of 10°/s, were presented in a grid of 7 × 14 positions, with the centers of the looming stimuli separated by ∼9° (Figure 3). The grid in Figure 3A represents the locations of the looming stimuli centers on the display, while the eye viewing transformation (Figure 3—figure supplement 1B) was used to position the stimulus responses more accurately in spherical coordinates from the fly eye perspective (Figure 3B, C).

The looming and looming related stimuli used throughout the study are listed in Table 2. Most stimuli progressed with an ‘r/v’ time course, detailed below. The details of the stimulus types listed in the table (first column) are as follows:

1. Small looming RF mapping stimulus: Maximal contrast dark disc, over mid-contrast grey background expanding from ∼4.5֯ to ∼18֯.
2. Dark Looming Disc: Maximal contrast dark disc, over mid-contrast grey background expanding from ∼4.5֯ to ∼54֯.
3. Bright Looming Disc: Maximal contrast bright looming disc stimulus, over mid-contrast grey background expanding from ∼4.5֯ to ∼54֯.
4. Dark Constant Looming Disc: Dark Looming Disc stimulus frames presented with constant speed profile.
5. Luminance-matched stimulus: ∼54֯ disc darkening with the same temporal profile as Dark Looming Disc stimulus. Intended to isolate the luminance component of the looming stimulus.
6. Looming annulus stimulus: Only the edge portion of the Dark Looming Disc stimulus. Intended to isolate the edge-motion component of the looming stimulus.
7. Bilateral Looming: Same as Dark Looming Disc but presented at symmetrical positions on both hemi fields of the visual display.
8. Small object motion stimulus: Maximal contrast dark ∼9֯ x∼9 ֯ small square over mid-contrast grey background, sweeping across display.
9. Vertical bar motion stimulus: Maximal contrast dark ∼9 ֯ width, full height vertical bar over mid-contrast grey background, sweeping across display.

To better match the receptive field of LC6G2 cells, the stimuli were positioned above the equator (18° elevation for looming stimuli, 31.5° elevation for the small object).

The stimuli were generated using custom MATLAB (Mathworks, Natick, MA) scripts (https://github.com/mmmorimoto/visual-stimuli). Schematics of the loom-related stimuli are shown in Figure 4B. The dark/bright loom stimuli each consisted of a series of 35 disk sizes, with the edge pixel intensity interpolated to approximate a circle on the discrete LED screen. The luminance-matched stimulus was created using the dark looming disk stimulus, spatially scrambling the location of dark pixels of each frame only within the area of the final size of the disk. The looming annulus was created by masking away the inner diameter (leaving the outer 9° diameter) of the dark disc stimulus for every frame. The time series of looming stimuli sizes were presented based on the classic parameterization for looming stimuli assuming a constant velocity of approach. The angular size of the object (θ) increases according to the equation θ(t) =2×tan^-1^(r/vt), where r is the radius of the object, v is its approach speed. Speed of the loom is represented by the ratio of these parameters (r/v, Figure 4B, tuning curves) (Gabbiani et al., 1999). The experimental protocol consisted of 3 repetitions of each stimulus type presented using a randomized block trial structure. Stimulus epochs were interleaved with at minimum 2 seconds of blank frame epochs that allowed the GCaMP6m fluorescence to decay back to baseline. Each protocol lasted 15-20mins and subsequently presented 3 times, resulting in the total experiment time of ∼ 1 hour.

### *In vivo* 2-photon calcium imaging: data analysis

Data were analyzed with software written in MATLAB. Motion correction was performed by cross-correlating each frame to a mean reference image and maximizing the correlation iteratively (using https://bitbucket.org/jastrother/neuron_image_analysis). The fluorescence signal was determined within hand-drawn regions of interest selected to tightly enclose the entire slice of each glomerulus captured within the imaging plane. ΔF/F was calculated as the ratio of (F - F0) / F0, where F is the instantaneous fluorescence signal and F0 is calculated as the 10^th^ percentile of the fluorescence signal (a robust estimator of baseline levels) within a sliding 300 frame window. These parameters were determined empirically to optimally fit the actual baseline fluorescence. For combining responses of individual flies across animals, we normalized the ΔF/F responses from each individual fly to the 98^th^ percentile of the ΔF/F across all visual stimuli within one experiment. The 98^th^ percentile (a robust estimator of peak levels) was typically near where the curve of the cumulative distribution of all pixel values in an experiment shows saturation. All grouped responses are the mean of the mean response (across repeated stimulus presentations) of all flies in the dataset. For LC6 responses in Figure 4, data from 4/6 flies were also used in our previous publication (Wu et al., 2016). Error bars for tuning curves and response curves in Figure 2,4 and 2—figure supplement 1 indicate mean ± SEM. All significance results presented for calcium imaging were determined with the Mann-Whitney U test. Pearson’s correlation coefficient (r) was used to assess the degree of correlation.

### EM Reconstruction

LC6s and their target neurons were manually reconstructed in a serial section transmission electron microscopy (ssTEM) volume of a full adult female *Drosophila melanogaster* brain (FAFB; Figure 5A) (Zheng et al., 2018) using CATMAID (Saalfeld et al., 2009). We followed established guidelines (Schneider-Mizell et al., 2016) for tracing skeletons for each neuron and identifying synaptic connections. We found and traced 65 LC6 neurons in the right hemisphere of the FAFB volume but visualized in Figure 5 and 6 as the left side. Surprisingly, the LC6\LC9 tract was shifted somewhat medially in this brain, a feature that has also occasionally been observed with light microscopy (http://flycircuit.tw). The morphology of each neuron’s dendrites was only approximately reconstructed, favoring completion of the largest branches but not completing the finest dendritic branches. We believe this is adequate for determining the retinotopic location of the center of each neuron’s anatomical receptive field, but the boundary of each receptive field might be underestimated. The axon terminals of each LC6 neuron were completely reconstructed and all identifiable synapses on the axons were tagged. The volume demarcated by all LC6 pre-synapses was used to generate a mask identifying the LC6 glomerulus. We then followed the LC6 synapses within the glomerulus to identify synaptic partners. By comparing the synaptic partners with the morphology of the expected target neurons from light microscopy data (Figure 1 and Figure 1—figure supplement 1), we identified the target neurons which we further reconstructed for the analysis in Figure 6. We reconstructed 2 LC6G1 bilateral neurons with cell bodies on the right-hand side and two with cell bodies on the left-hand side. We initially reconstructed 2 of the ipsilateral LC6G1 neurons as well, but after finding variable receptive field structures we searched extensively and found 3 more neurons of this same morphology.

All target neurons’ morphologies (except for the left-hand-side LC6G1 neurons) as well as the connections between the target neurons and LC6 neurons within the right-hand side glomerulus were traced to completion and reviewed by an independent team member.

### Computational analysis of EM reconstructions

All EM analyses and visualizations were carried out in the R language (R Core Team, 2013) using open-source packages, mainly the “natverse” (http://natverse.org/) (Bates et al., 2019). In order to estimate the retinotopic correspondence for each reconstructed LC6 neuron we mapped the visual field of one eye onto a layer of the lobula. We fit a 2^nd^ order curved surface through the dendrites of all LC6 neurons (Figure 5B, C). In an ongoing effort we are reconstructing multiple neurons types throughout the medulla to precisely identify the location of every column in the right optic lobe. While this effort is beyond the scope of the work we describe in this manuscript, we used these data to identify two medulla columns that correspond to the center of the eye and a position on the central meridian (the line that partitions the eye between anterior and posterior halves). We expect these points are within 1- 2 columns of these ideal locations. We then identified columnar neurons that connect to LC6 neurons and found one subtype of Tm5 neurons (Karuppudurai et al., 2014) that are highly presynaptic to LC6 dendrites in the lobula. We reconstructed Tm5 neurons from the two medulla columns, extending these reference points from the medulla to the surface defined by the LC6 dendrites in the lobula (Figure 5B, C). This allowed us to map the curved surface onto a hemisphere. Or more precisely, a spherical lune with [-90°, +90°] in longitude and [−10°, +160°] in latitude: an approximation of the field of view of one eye (see eye mapping in Figure 3—figure supplement 1, and description above). Consequently, we mapped each LC6 neuron’s location (center-of-mass of the dendrite) onto eye coordinates. Figure 5D shows the central locations and the boundaries of all LC6 neurons projected onto a flattened field of view of one eye.

We found over 1900 synaptic connections between LC6 neurons in the glomerulus. We tested whether these connections were distributed at random or were more likely to occur between neurons with nearby visual coordinates. For all possible LC6 pairs, we multiplied the distances between dendrite centers in the lobula by the corresponding number of LC6-LC6 synapses in the glomerulus, and then calculated the mean value over all 64 LC6-LC6 pairs for a given LC6 (Figure 5E). As a comparison we calculated the weighted mean after randomly shuffling these 64 distances.

To assess whether the glomerulus exhibited some retinotopic organization that was not visible at the light level, we divided the glomerulus in 10 compartments along its long axis, each containing 1/10 of LC6 post-synapses. Since synaptic connections are often polyadic (mostly one-to-many) in fly brains, we used the single presynaptic connectors and ignored the possible numerous postsynaptic connectors for dividing up the glomerulus. For each compartment, we constructed a 2D multi-Gaussian (65 Gaussians for 65 LC6 neurons) distribution in the eye coordinate (Figure 5F). The center of each LC6 Gaussian was its dendritic center. Its height was proportional to the synapse count in the given compartment.

The half-widths of all Gaussians were the same and assigned as the radius of the dendritic arbors averaged over all 65 LC6 neurons (15.5 µm). The 65 Gaussian distributions are then summed to produce a 2D multi-Gaussian distribution for each compartment.

Similarly, we also constructed 2D multi-Gaussian distributions for LC6 target neurons. Here the centers of the Gaussians were again the dendritic centers of the LC6s while their heights were the synapse count for a given target neuron. This procedure results in an estimate of the feed-forward, anatomical receptive field, based only on neuron morphology and connectivity. To compare this prediction to *in vivo* imaging data we have summed the anatomical receptive field estimates from groups of individual neurons to mimic the summed responses from genetically encoded calcium indicator expressed in populations of neurons of the same type (Figure 6 C). As these data have been transformed to spherical coordinates in an eye-centered reference frame, they are directly aligned to the receptive fields measured using calcium imaging based on the eye projection (Figure 3—figure supplement 1). The composite images are shown in Figure 6D). All reconstructed neurons described in the manuscript will be available at https://fafb.catmaid.virtualflybrain.org/, and all data analysis code is available at https://github.com/reiserlab/LC6downstream.

## Acknowledgements

We thank the Janelia Fly Light Project Team for help with imaging driver lines and processing of FISH samples. We thank Jasmine Le for training and assistance with the functional connectivity experiments. We thank members of the Bock lab, especially Scott Lauritzen, for supporting our early EM reconstruction efforts and Gregory Jefferis for introducing us to his tools for computational neuroanatomy. We thank Janelia’s Connectome Annotators: Padideh Ghorbani, N. Aidan Smith, Marisa Dreher, Miriam Flynn, Connor Laughland, Henrique Ludwig, Alex Thomson, Bruck Gezahegn, supervised by Ruchi Parekh, for their dedicated reconstruction of the neuronal circuits featured in this study. We are also grateful to members of the Reiser Lab, especially Eyal Gruntman, Frank Loesche, and Kit Longden, for comments on the manuscript. This project was supported by HHMI.

## Author Contributions

Conceptualization, M.M.M. and M.B.R.; Methodology, M.M.M., A.M.W., M.D.I., D.D.B., and M.B.R.; Software, M.M.M., A.Z., A.M.W., and M.D.I.; Formal Analysis, M.M.M., A.N., and A.Z.; Investigation, M.M.M. and A.N.; Resources, A.N., E.M.R., and D.D.B.; Writing – Original Draft, M.M.M., A.N., A.Z., and M.B.R.; Writing – Review & Editing, M.M.M., A.N., A.Z., D.D.B., G.M.R., and M.B.R.; Supervision: M.B.R.; Funding Acquisition: D.D.B., G.M.R., and M.B.R.

---

**Figure 1—figure supplement 1:**
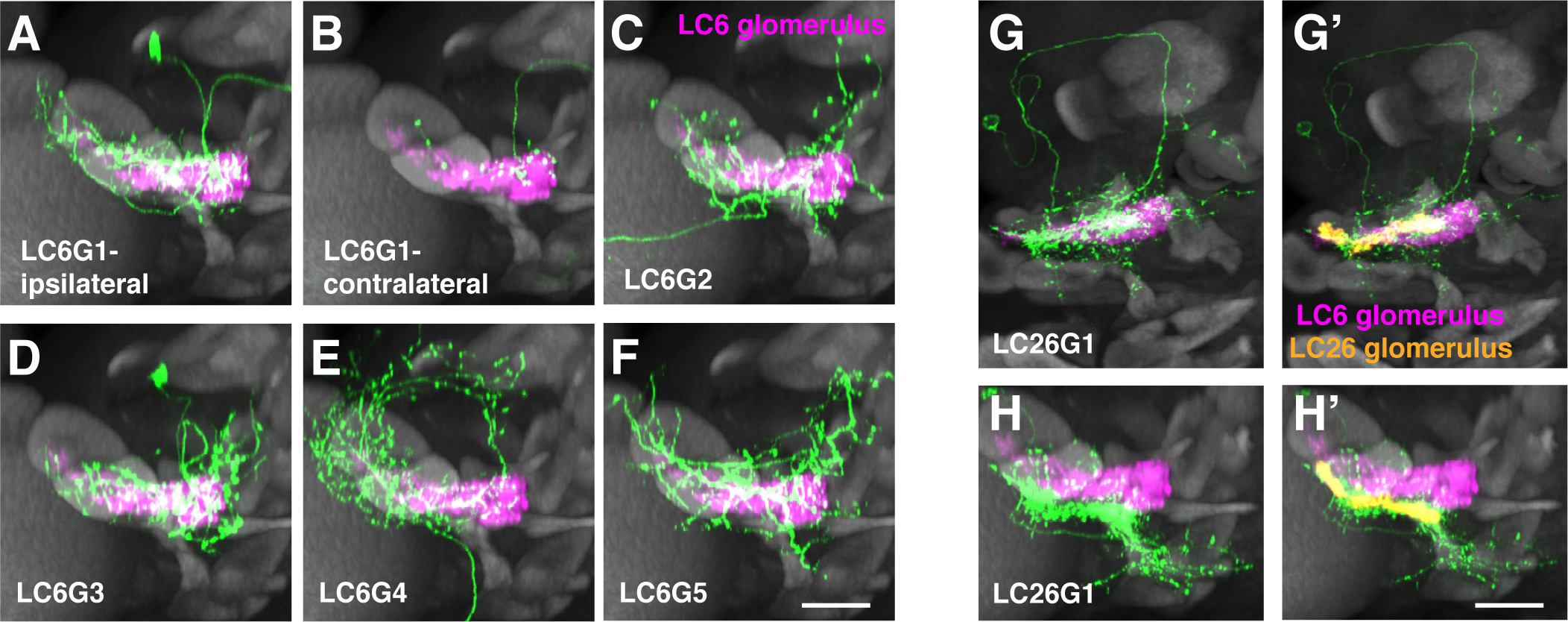
Additional anatomical details of LC6G neurons. **(A-F)** Views of the cells shown in Figure 1H-M rotated by 90° around the mediolateral axis. **(G-H)** Segmented LC26G1 cells displayed in the same views as the LC6G cells in Figure 1 **H-M (G, G’)** and Figure 1—figure supplement 1**A-F (H, H’)**. The LC26 glomerulus outline is based on syt-HA expression in LC26 driven by an LC26 split-GAL4 driver (Wu et al., 2016). Note that LC26G1 shows only minor overlap with the LC6 glomerulus. Scale bar, 20 µm.

**Figure 1—figure supplement 2:**
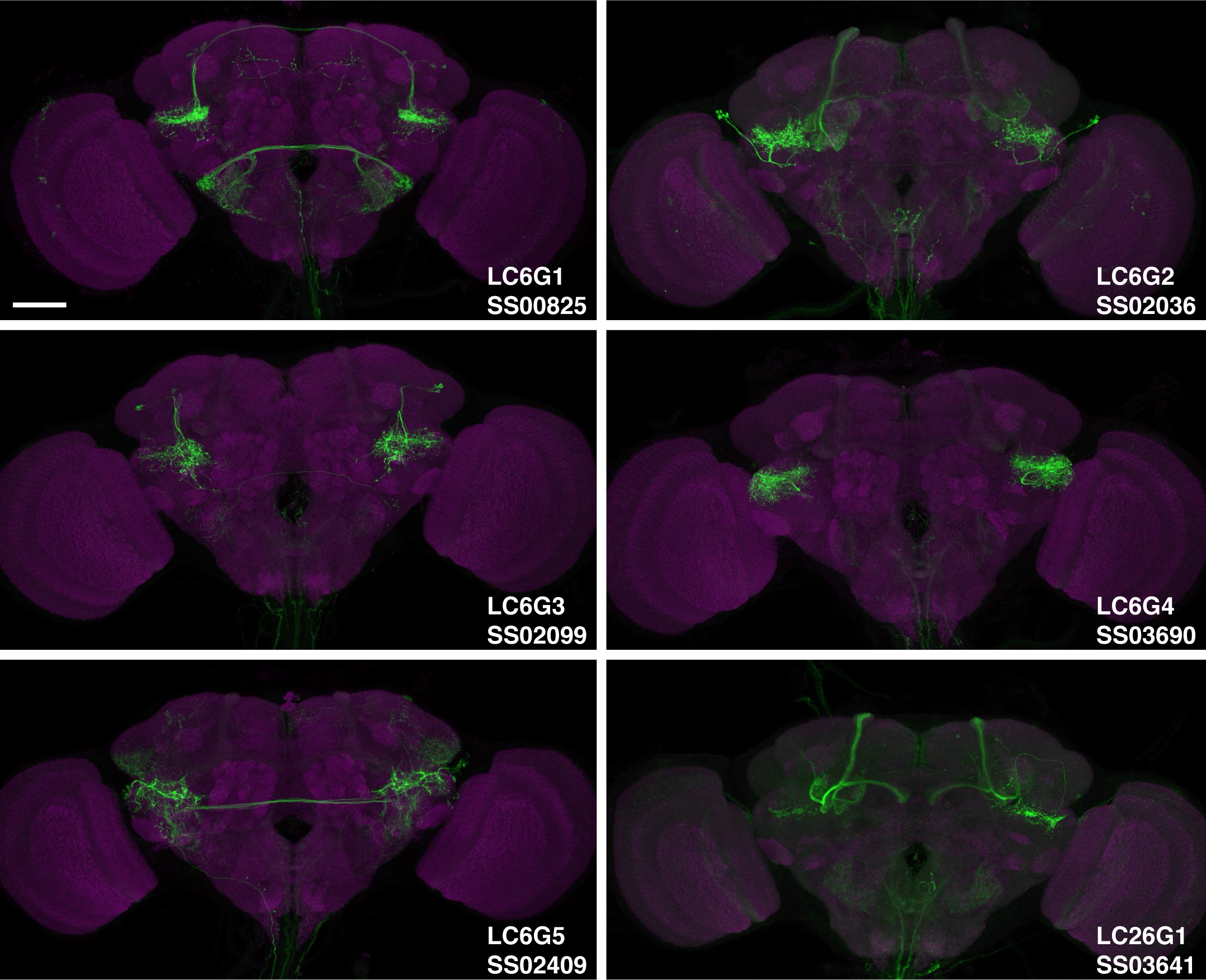
Expression patterns of LC6G split-GAL4 lines. Maximum intensity projections through confocal stacks of brains immunolabeled for a GAL4 driven membrane marker (green) and a reference label (anti-Brp, magenta). Imaging parameters and post-imaging adjustments of brightness and contrast are not identical for different images. Scale bar, 50 µm. The line name and expressed cell type for each split-GAL4 line is indicated in the lower right corner. See Table 1 for detailed summary of driver lines used in this study.

**Figure 1—figure supplement 3:**
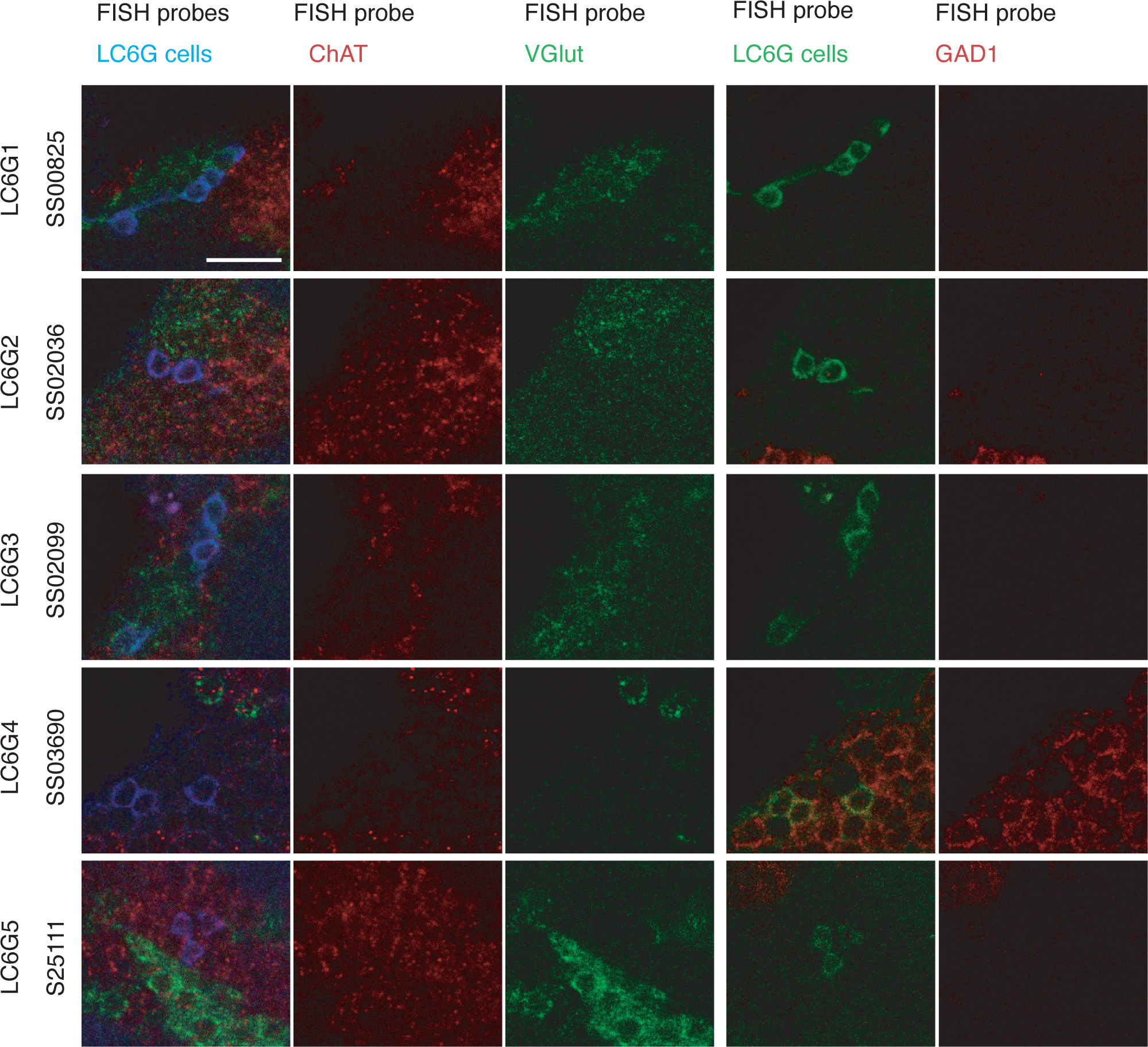
FISH detection of neurotransmitter markers in LC6G neurons. **(A)** Transcripts of GAD1, VGlut and ChaT were detected by FISH as described in (Meissner et al., 2019). Images show single slices of confocal stacks. Images for each driver line show the same group of cells but were acquired in two sets. The 1^st^ and 4^th^ columns are composites with the colors labeled along the top row indicating each probe. Cell bodies of cells with the same neurotransmitter phenotype are often found in groups (presumably from the same lineage). For example, both LC6G1 and LC6G3 cells are part of the same cluster of apparently glutamatergic neurons. Scale bar, 10 µm.

**Figure 2—figure supplement 1:**
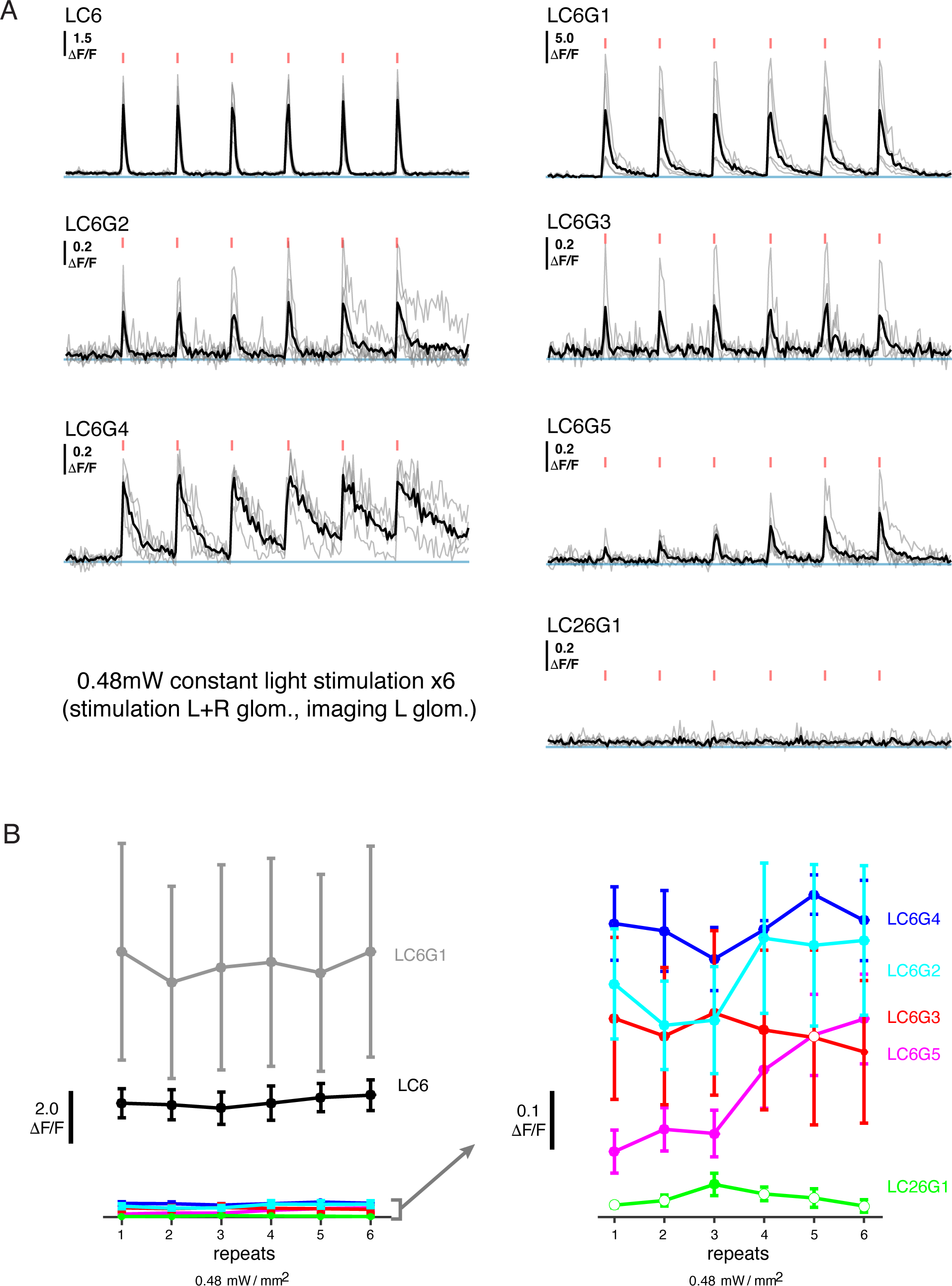
Additional functional connectivity results with alternative stimulation protocol and an additional cell type. **(A)** Calcium responses of candidate downstream neuron types in response to LC6 Chrimson activation (N=4-5 per combination; same flies as in Figure 2; individual mean sample response in gray, mean response across flies in black). Red tick marks indicate the activation stimulus. In this protocol, a constant illumination level was used for each stimulation repeat. **(B)** Peak response means (± SEM) are shown for each candidate downstream cell-type. Closed circles denote data points significantly different from pre-stimulus baseline, while open circles denote data points not significantly different from pre-stimulus baseline (p<0.1, two sample T-test). See Table 1 for detailed summary of driver lines used in this study.

**Figure 3—figure supplement 1:**
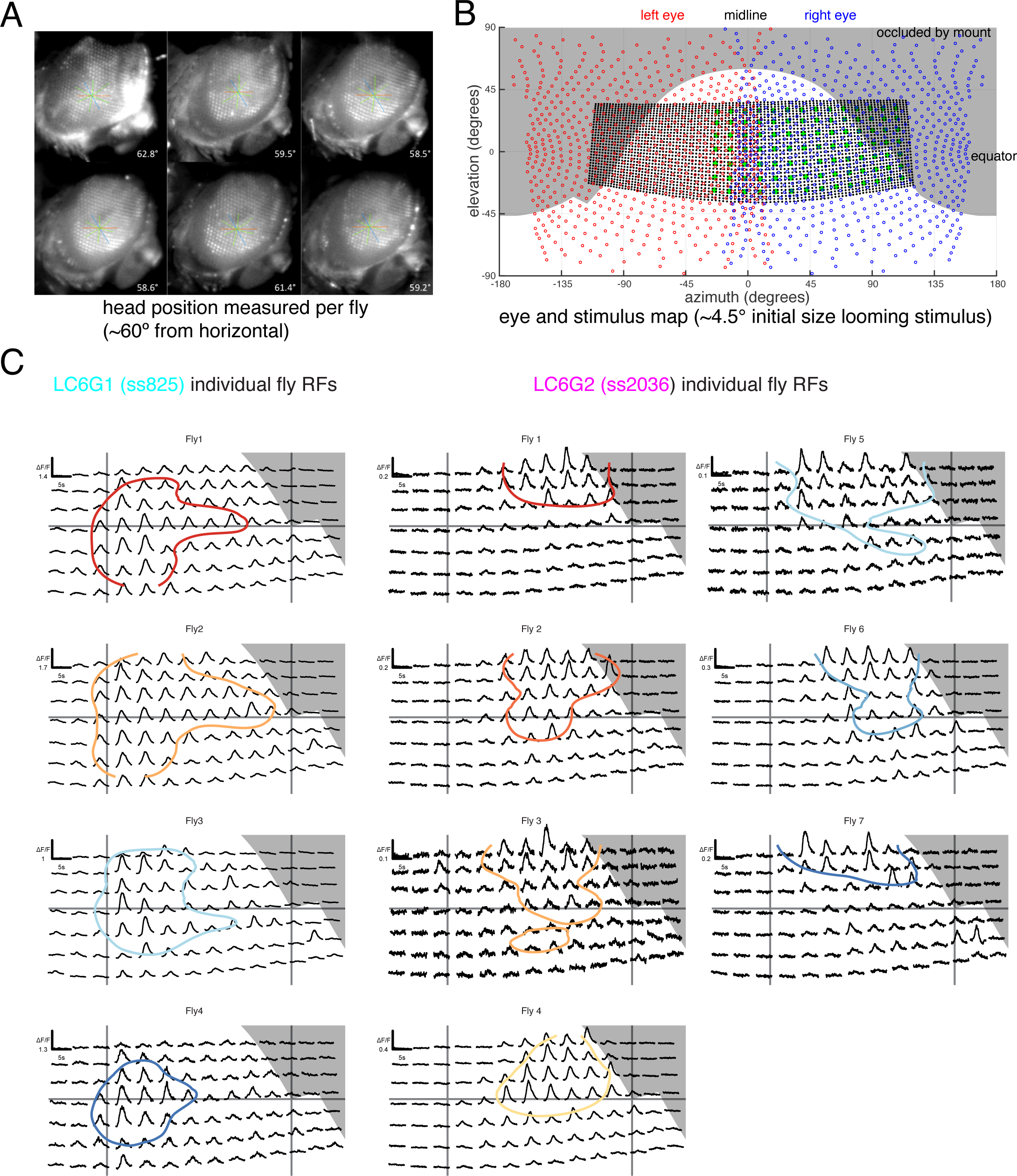
Eye map and individual fly receptive fields. **(A)** Side view of head fixed fly. The ommatidial axes were used to determine the inclination of the head. Great care was taken to consistently align the fly head across preparations. **(B)** Overlay of ommatidial and stimulus maps in visual space coordinates. The extent of visual stimulation panels is shown as the black dots, each representing a single LED of the 32 × 96 array, while the green disc represents the center positions of each small looming stimulus used for RF mapping. Grey background areas were estimated to not be visible to the fly, from occlusion by head mount. **(C)** Calcium responses of LC6G1 and LC6G2 neurons, measured from individual flies, to the RF mapping stimuli (Figure 3A) Plotting conventions follow Figure 3B, and the colors for individual flies are carried over from Figure 3C. The accuracy of our alignment between individual flies and our visual stimulus is confirmed by the lack of visual responses for the occluded positions in the upper right corner (in grey). See Table 1 for detailed summary of driver lines used in this study.

**Figure 5—figure supplement 1:**
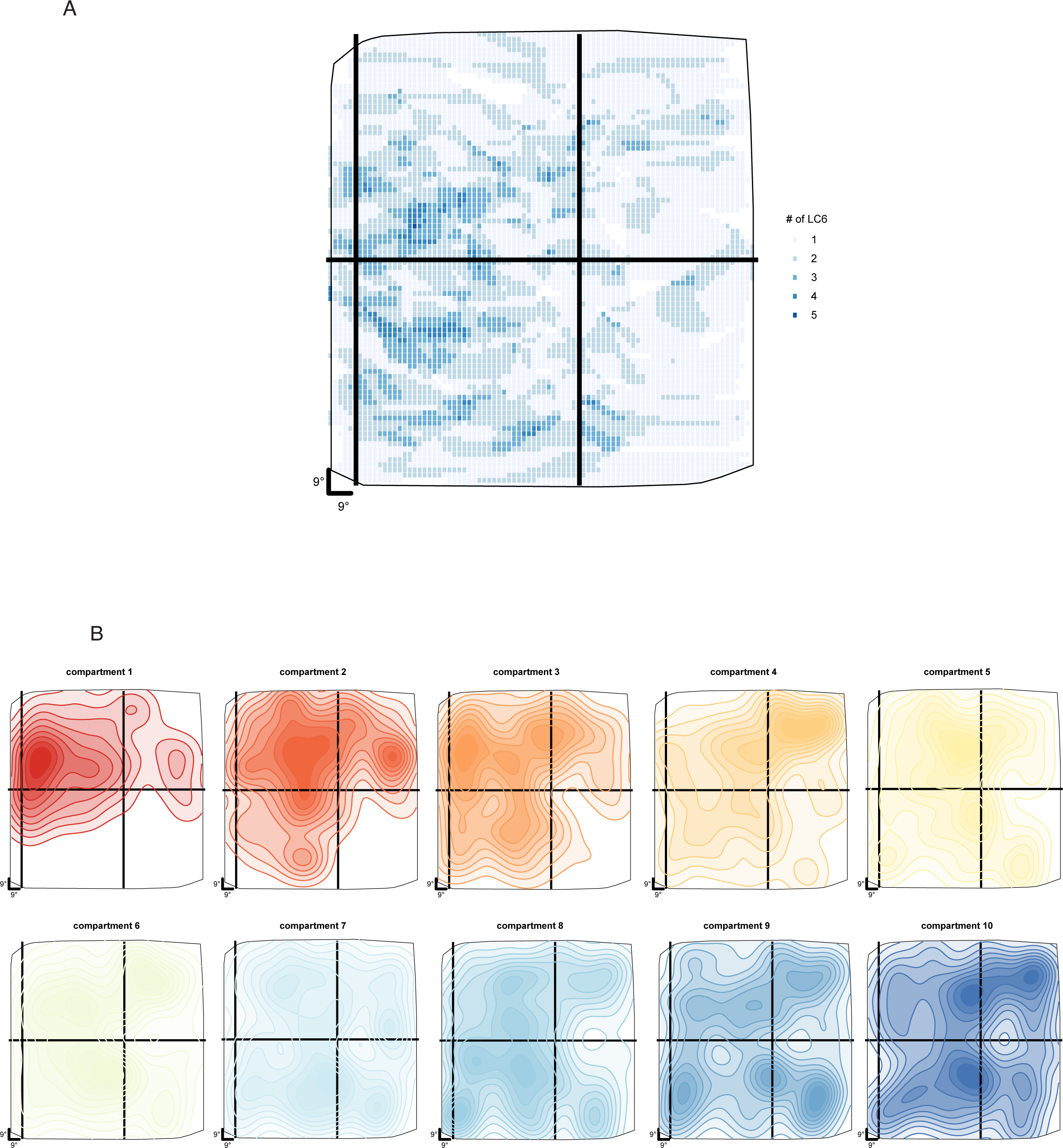
Distribution of LC6 arbors in the lobula from EM data. **(A)** Coverage of the eye by LC6 dendrites based on the estimated anatomical receptive fields of Figure 5D. The numbers indicate the number of LC6 neurons with overlapping receptive fields at each sampled location within this projection of the lobula layer 4. The plotting convention follows Figure 5D. **(B)** Contour plots of the LC6-LC6 synapse-weighted spatial receptive fields of each of 10 LC6 glomerulus compartments (Figure 5F).

**Figure 6—figure supplement 1:**
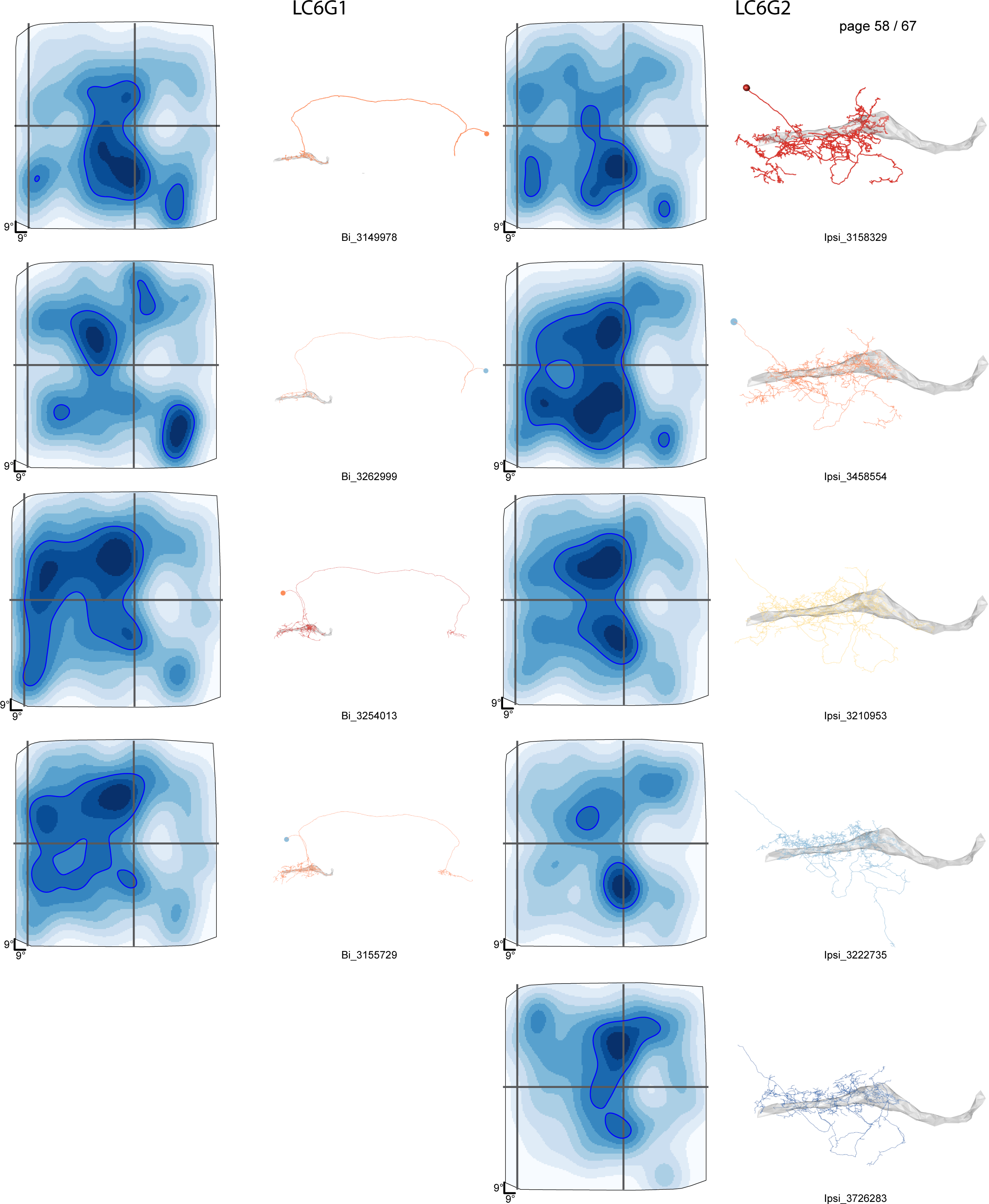
Details of reconstructed LC6G neuron morphology and estimated anatomical Receptive Fields. The names for each reconstructed neurons match the labels/annotation for each skeleton (to be posted at https://fafb.catmaid.virtualflybrain.org/).

**Figure 6—figure supplement 2:**
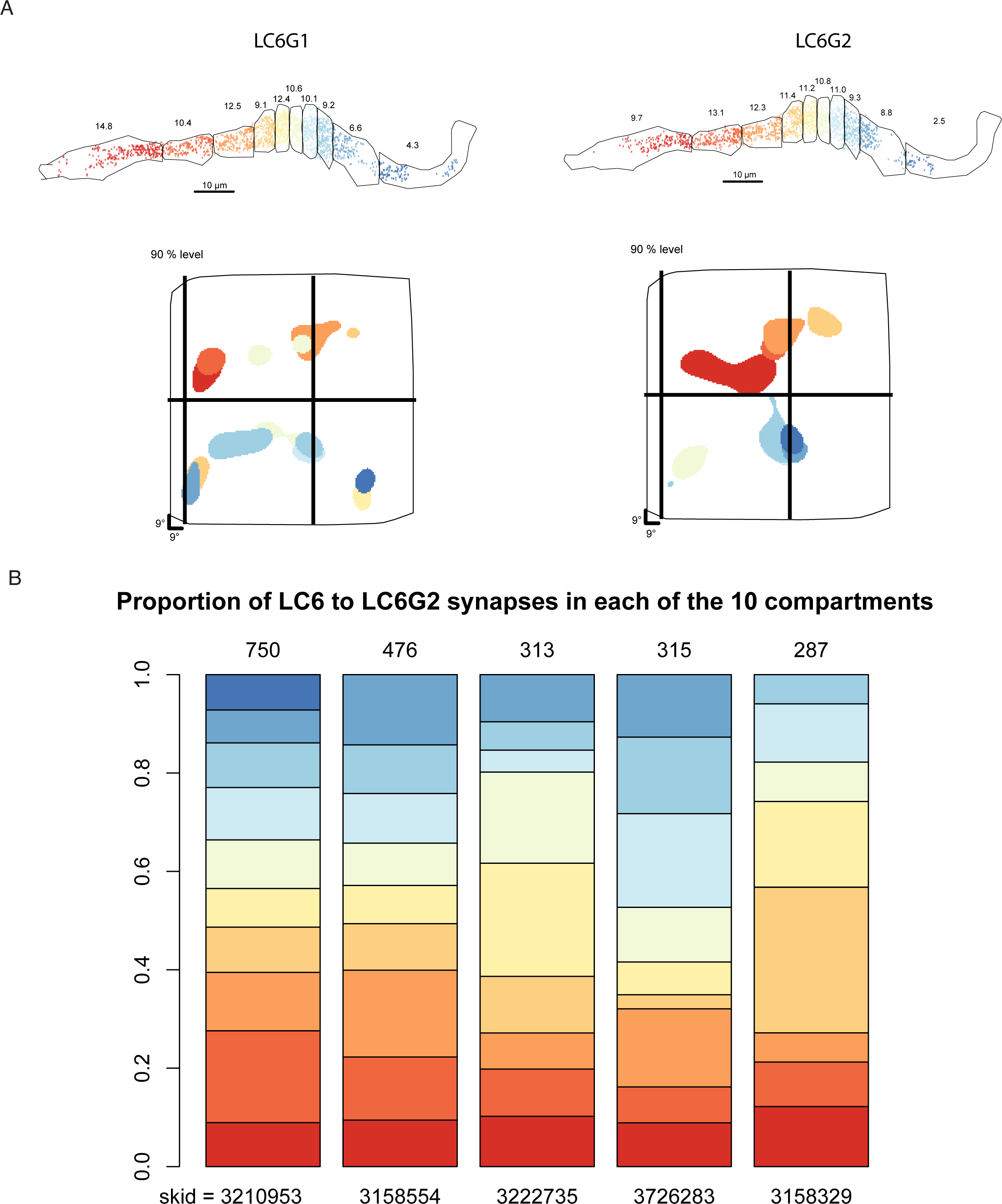
Anatomical Receptive Field analysis in 10 compartments of the LC6 glomerulus. **(A)** The peak RF for the LC6 inputs from each compartment of the glomerulus, same analysis as for Figure 5F, but here only plotting the 90% level. The LC6G1 and LC6G2 inputs, combined across all cells used in the Figure 6B analysis, are shown. **(B)** A comparison of the distribution of LC6 inputs to the 5 LC6G2 neurons throughout the glomerulus. The 10 compartments are defined using all LC6 pre-synapses (as in Figure 5F). The listed names are the numerical component of the Skeleton ID for each reconstructed neuron and match the names in Figure 6—figure supplement 1.

## References

1. Ache JM, Polsky J, Alghailani S, Parekh R, Breads P, Peek MY, Bock DD, von Reyn CR, Card GM. 2019. Neural Basis for Looming Size and Velocity Encoding in the Drosophila Giant Fiber Escape Pathway. Curr Biol 29:1073–1081.e4. doi:10.1016/J.CUB.2019.01.079

2. Aso Y, Hattori D, Yu Y, Johnston RM, Iyer NA, Ngo TTB, Dionne H, Abbott LF, Axel R, Tanimoto H, Rubin GM. 2014. The neuronal architecture of the mushroom body provides a logic for associative learning. Elife. doi:10.7554/eLife.04577

3. Bates AS, Manton JD, Jagannathan SR, Costa M, Schlegel P, Rohlfing T, Jefferis GSXE. 2019. The natverse: a versatile computational toolbox to combine and analyse neuroanatomical data. bioRxiv 6353. doi:10.1101/006353

4. Benjamini Y, Hochberg Y. 1995. Controlling the False Discovery Rate: A Practical and Powerful Approach to Multiple Testing. J R Stat Soc Ser B. doi:10.1111/j.2517-6161.1995.tb02031.x

5. Bogovic JA, Otsuna H, Heinrich L, Ito M, Jeter J, Meissner G, Nern A, Colonell J, Malkesman O, Ito K, Saalfeld S. 2018. An unbiased template of the Drosophila brain and ventral nerve cord. bioRxiv. doi:10.1101/376384

6. Buchner E. 1971. Dunkelanregung des stationaeren Flugs der Fruchtfliege Drosophila. University of Tuebingen.

7. Card G, Dickinson MH. 2008. Visually Mediated Motor Planning in the Escape Response of Drosophila. Curr Biol 18:1300–1307. doi:10.1016/j.cub.2008.07.094

8. Chen T-W, Wardill TJ, Sun Y, Pulver SR, Renninger SL, Baohan A, Schreiter ER, Kerr RA, Orger MB, Jayaraman V, Looger LL, Svoboda K, Kim DS. 2013. Ultrasensitive fluorescent proteins for imaging neuronal activity. Nature 499:295–300. doi:10.1038/nature12354

9. Chen T, Wardill TJ, Sun Y, Pulver SR, Renninger SL, Baohan A, Schreiter ER, Kerr RA, Orger MB, Jayaraman V, Looger LL, Svoboda K, Kim DS. 2013. Ultrasensitive fluorescent proteins for imaging neuronal activity. Nature 499:295–300. doi:10.1038/nature12354

10. Chklovskii DB, Koulakov AA. 2004. MAPS IN THE BRAIN: What Can We Learn from Them? Annu Rev Neurosci. doi:10.1146/annurev.neuro.27.070203.144226

11. Davis FP, Nern A, Picard S, Reiser MB, Rubin GM, Eddy SR, Henry GL. 2020. A genetic, genomic, and computational resource for exploring neural circuit function. Elife. doi:10.7554/elife.50901

12. Demerec M. 1965. Biology of Drosophila. New York: Hafner Press.

13. Dionne H, Hibbard KL, Cavallaro A, Kao JC, Rubin GM. 2018. Genetic reagents for making split-GAL4 lines in Drosophila. Genetics. doi:10.1534/genetics.118.300682

14. Fischbach K-F, Dittrich APM. 1989. The optic lobe of Drosophila melanogaster. I. A Golgi analysis of wild-type structure. Cell Tissue Res 258. doi:10.1007/BF00218858

15. Freeman J, Vladimirov N, Kawashima T, Mu Y, Sofroniew NJ, Bennett D V, Rosen J, Yang C-T, Looger LL, Ahrens MB. 2014. Mapping brain activity at scale with cluster computing. Nat Methods 11:941–950. doi:10.1038/nmeth.3041

16. Gabbiani F, Krapp HG, Laurent G. 1999. Computation of object approach by a wide-field, motion-sensitive neuron. J Neurosci 19:1122–41. doi:10.1523/JNEUROSCI.19-03-01122.1999

17. Garrett ME, Nauhaus I, Marshel JH, Callaway EM, Garrett ME, Marshel JH, Nauhaus I, Garrett ME. 2014. Topography and areal organization of mouse visual cortex. J Neurosci. doi:10.1523/JNEUROSCI.1124-14.2014

18. Hubel DH, Wiesel TN. 1962. Receptive fields, binocular interaction and functional architecture in the cat’s visual cortex. J Physiol 160:106–154. doi:10.1113/jphysiol.1962.sp006837

19. Jenett A, Rubin GM, Ngo T-TB, Shepherd D, Murphy C, Dionne H, Pfeiffer BD, Cavallaro A, Hall D, Jeter J, Iyer N, Fetter D, Hausenfluck JH, Peng H, Trautman ET, Svirskas RR, Myers EW, Iwinski ZR, Aso Y, DePasquale GM, Enos A, Hulamm P, Lam SCB, Li H-H, Laverty TR, Long F, Qu L, Murphy SD, Rokicki K, Safford T, Shaw K, Simpson JH, Sowell A, Tae S, Yu Y, Zugates CT. 2012a. A GAL4-Driver Line Resource for Drosophila Neurobiology. Cell Rep 2:991–1001. doi:10.1016/j.celrep.2012.09.011

20. Jenett A, Rubin GM, Ngo T-TB, Shepherd D, Murphy C, Dionne H, Pfeiffer BD, Cavallaro A, Hall D, Jeter J, Iyer N, Fetter D, Hausenfluck JH, Peng H, Trautman ET, Svirskas RR, Myers EW, Iwinski ZR, Aso Y, DePasquale GM, Enos A, Hulamm P, Lam SCB, Li H-H, Laverty TR, Long F, Qu L, Murphy SD, Rokicki K, Safford T, Shaw K, Simpson JH, Sowell A, Tae S, Yu Y, Zugates CT. 2012b. A GAL4-Driver Line Resource for Drosophila Neurobiology. Cell Rep 2:991–1001. doi:10.1016/j.celrep.2012.09.011

21. Karuppudurai T, Lin TY, Ting CY, Pursley R, Melnattur K V., Diao F, White BH, Macpherson LJ, Gallio M, Pohida T, Lee CH. 2014. A Hard-Wired Glutamatergic Circuit Pools and Relays UV Signals to Mediate Spectral Preference in Drosophila. Neuron. doi:10.1016/j.neuron.2013.12.010

22. Keleş MF, Frye MA. 2017. Object-Detecting Neurons in Drosophila. Curr Biol 27:680–687. doi:10.1016/j.cub.2017.01.012

23. Klapoetke NC, Murata Y, Kim SS, Pulver SR, Birdsey-Benson A, Cho YK, Morimoto TK, Chuong AS, Carpenter EJ, Tian Z, Wang J, Xie Y, Yan Z, Zhang Y, Chow BY, Surek B, Melkonian M, Jayaraman V, Constantine-Paton M, Wong GK-S, Boyden ES. 2014. Independent optical excitation of distinct neural populations. Nat Methods 11:338–346. doi:10.1038/nmeth.2836

24. Klapoetke NC, Nern A, Peek MY, Rogers EM, Breads P, Rubin GM, Reiser MB, Card GM. 2017. Ultra-selective looming detection from radial motion opponency. Nature 551. doi:10.1038/nature24626

25. Knapp J-M, Chung P, Simpson JH. 2015. Generating Customized Transgene Landing Sites and Multi-Transgene Arrays in *Drosophila* Using phiC31 Integrase. Genetics 199:919– 934. doi:10.1534/genetics.114.173187

26. Kvon EZ, Kazmar T, Stampfel G, Yáñez-Cuna JO, Pagani M, Schernhuber K, Dickson BJ, Stark A. 2014. Genome-scale functional characterization of Drosophila developmental enhancers in vivo. Nature 512:91–95. doi:10.1038/nature13395

27. Lai S-L, Lee T. 2006. Genetic mosaic with dual binary transcriptional systems in Drosophila. Nat Neurosci 9:703–709. doi:10.1038/nn1681

28. Liu WW, Wilson RI. 2013. Glutamate is an inhibitory neurotransmitter in the Drosophila olfactory system. Proc Natl Acad Sci U S A. doi:10.1073/pnas.1220560110

29. Luan H, Peabody NC, Vinson CR, White BH. 2006. Refined spatial manipulation of neuronal function by combinatorial restriction of transgene expression. Neuron 52:425–436.

30. Mauss AS, Pankova K, Arenz A, Nern A, Rubin GM, Borst A. 2015. Neural Circuit to Integrate Opposing Motions in the Visual Field. Cell. doi:10.1016/j.cell.2015.06.035

31. Meissner GW, Aljoschanern, Singer RH, Wong AM, Malkesman O, Long X. 2019. Mapping neurotransmitter identity in the whole-mount drosophila brain using multiplex high-throughput fluorescence in situ hybridization. Genetics. doi:10.1534/genetics.118.301749

32. Mu L, Ito K, Bacon JP, Strausfeld NJ. 2012. Optic Glomeruli and Their Inputs in Drosophila Share an Organizational Ground Pattern with the Antennal Lobes. J Neurosci 32:6061– 6071. doi:10.1523/JNEUROSCI.0221-12.2012

33. Namiki S, Dickinson MH, Wong AM, Korff W, Card GM. 2018. The functional organization of descending sensory-motor pathways in Drosophila. Elife 7. doi:10.7554/eLife.34272

34. Nern A, Pfeiffer BD, Rubin GM. 2015. Optimized tools for multicolor stochastic labeling reveal diverse stereotyped cell arrangements in the fly visual system. Proc Natl Acad Sci 112:E2967–E2976. doi:10.1073/pnas.1506763112

35. Otsuna H, Ito K. 2006. Systematic analysis of the visual projection neurons of Drosophila melanogaster. I. Lobula-specific pathways. J Comp Neurol. doi:10.1002/cne.21015

36. Panser K, Tirian L, Schulze F, Villalba S, Jefferis GSXE, Bühler K, Straw AD. 2016. Automatic Segmentation of Drosophila Neural Compartments Using GAL4 Expression Data Reveals Novel Visual Pathways. Curr Biol 26:1943–1954. doi:10.1016/j.cub.2016.05.052

37. Pfeiffer BD, Ngo T-TB, Hibbard KL, Murphy C, Jenett A, Truman JW, Rubin GM. 2010. Refinement of Tools for Targeted Gene Expression in Drosophila. Genetics 186:735– 755. doi:10.1534/genetics.110.119917

38. Reiser MB, Dickinson MH. 2008. A modular display system for insect behavioral neuroscience. J Neurosci Methods 167:127–139. doi:10.1016/j.jneumeth.2007.07.019

39. Ribeiro IMA, Drews M, Bahl A, Machacek C, Borst A, Dickson BJ. 2018. Visual Projection Neurons Mediating Directed Courtship in Drosophila. Cell 174:607–621.e18. doi:10.1016/j.cell.2018.06.020

40. Saalfeld S, Cardona A, Hartenstein V, Tomančák P. 2009. CATMAID: Collaborative annotation toolkit for massive amounts of image data. Bioinformatics. doi:10.1093/bioinformatics/btp266

41. Schneider-Mizell CM, Gerhard S, Longair M, Kazimiers T, Li F, Zwart MF, Champion A, Midgley FM, Fetter RD, Saalfeld S, Cardona A. 2016. Quantitative neuroanatomy for connectomics in Drosophila. Elife. doi:10.7554/eLife.12059

42. Sen R, Wu M, Branson K, Robie A, Rubin GM, Dickson BJ. 2017. Moonwalker Descending Neurons Mediate Visually Evoked Retreat in Drosophila. Curr Biol 27:766–771. doi:10.1016/j.cub.2017.02.008

43. Shih CT, Sporns O, Yuan SL, Su TS, Lin YJ, Chuang CC, Wang TY, Lo CC, Greenspan RJ, Chiang AS. 2015. Connectomics-based analysis of information flow in the drosophila brain. Curr Biol. doi:10.1016/j.cub.2015.03.021

44. Shinomiya K, Huang G, Lu Z, Parag T, Xu CS, Aniceto R, Ansari N, Cheatham N, Lauchie S, Neace E, Ogundeyi O, Ordish C, Peel D, Shinomiya A, Smith C, Takemura S, Talebi I, Rivlin PK, Nern A, Scheffer LK, Plaza SM, Meinertzhagen IA. 2019. Comparisons between the ON- and OFF-edge motion pathways in the Drosophila brain. Elife. doi:10.7554/eLife.40025

45. Strausfeld NJ, Sinakevitch I, Okamura JY. 2007. Organization of local interneurons in optic glomeruli of the dipterous visual system and comparisons with the antennal lobes. Dev Neurobiol 67:1267–1288. doi:10.1002/dneu.20396

46. Strother JA, Nern A, Reiser MB. 2014. Direct observation of ON and OFF pathways in the Drosophila visual system. Curr Biol 24:976–83. doi:10.1016/j.cub.2014.03.017

47. Strother JA, Wu S-T, Wong AM, Nern A, Rogers EM, Le JQ, Rubin GM, Reiser MB. 2017. The Emergence of Directional Selectivity in the Visual Motion Pathway of Drosophila. Neuron 94:168–182.e10. doi:10.1016/j.neuron.2017.03.010

48. Sun Y, Nern A, Franconville R, Dana H, Schreiter ER, Looger LL, Svoboda K, Kim DS, Hermundstad AM, Jayaraman V. 2017. Neural signatures of dynamic stimulus selection in Drosophila. Nat Neurosci 20:1104–1113. doi:10.1038/nn.4581

49. Suver MP, Matheson AMM, Sarkar S, Damiata M, Schoppik D, Nagel KI. 2019. Encoding of Wind Direction by Central Neurons in Drosophila. Neuron 102:828–842.e7. doi:10.1016/j.neuron.2019.03.012

50. Tirian L, Dickson BJ. 2017. The VT GAL4, LexA, and split-GAL4 driver line collections for targeted expression in the Drosophila nervous system. bioRxiv. doi:10.1101/198648

51. Tootell RBH, Switkes E, Silverman MS, Hamilton SL. 1988. Functional anatomy of macaque striate cortex. II Retinotopic organization. J Neurosci. doi:10.1523/jneurosci.08-05-01531.1988

52. Wilson RI, Laurent G. 2005. Journal of Neuroscience. J Neurosci 23:4625–4634. doi:10.1523/jneurosci.2070-05.2005

53. Wu M, Nern A, Ryan Williamson W, Morimoto MM, Reiser MB, Card GM, Rubin GM. 2016. Visual projection neurons in the Drosophila lobula link feature detection to distinct behavioral programs. Elife 5. doi:10.7554/eLife.21022

54. Zheng Z, Lauritzen JS, Perlman E, Robinson CG, Nichols M, Milkie D, Torrens O, Price J, Fisher CB, Sharifi N, Calle-Schuler SA, Kmecova L, Ali IJ, Karsh B, Trautman ET, Bogovic JA, Hanslovsky P, Jefferis GSXE, Kazhdan M, Khairy K, Saalfeld S, Fetter RD, Bock DD. 2018. A Complete Electron Microscopy Volume of the Brain of Adult Drosophila melanogaster. Cell 174:730–743.e22. doi:10.1016/j.cell.2018.06.019

